# Apparent food selectivity reflects multiple non-food image properties

**DOI:** 10.64898/2026.09.18.751020

**Authors:** Cyn Fang, Meenakshi Khosla, Moshe Poliak, Nancy Kanwisher

## Abstract

Three recent publications have reported selective fMRI responses to images of food in the human ventral visual pathway. However, all three studies were based primarily on the Natural Scenes Dataset (NSD), in which high-level categories like food are correlated with other image properties such as the color and size of objects, and the distance of the scene. To test whether the reported food selectivity might reflect these or other correlates of food images rather than (or in addition to) food, we constructed novel stimuli that manipulated these properties on both food and non-food images, and collected data from new subjects in a data-rich design across two experiments pre-registered in OSF. We used a localizer paradigm based on a subset of NSD images to infer the “food component” in new subjects and then measured responses of this component to our new stimuli. In Experiment 1, we found that the food component response magnitude i) had a higher response to color than to greyscale images but showed no interaction between food and color, ii) showed a reduced preference for food over non-food when both were within reaching distance, and most importantly iii) was no higher for food than non-food when both were visually matched and presented as Cutouts on a white background. In Experiment 2, we found that the response of the food component could not be explained by object distance, real-world size, or mid-level visual image statistics. However, the response to non-food images with “gooey” material properties was as high as the response to food. Across both experiments, we consistently found a very low response (at or below fixation baseline) to NSD non-food images, which were predominantly outdoor scenes. Together, our results prompt a revision of prior claims including our own, indicating that the previously reported food component is better characterized as food-biased rather than strictly food-selective, and is driven in part by contextual or material features that are also present in non-food images. Our findings further highlight the importance of supplementing studies based on naturalistic images with experiments that unconfound image properties with carefully designed stimuli.

## Introduction

Few behaviors are more crucial to human survival than the ability to find and identify food. Although many sensory modalities contribute to locating and evaluating food, vision plays a central role. Given that food is such an important category of visual stimulus for humans, we might expect a neural population that responds selectively to images of food, as found for faces, places, and bodies (Kanwisher, 2025). Indeed, three recent papers have reported food selective responses in the human visual cortex (Khosla et al., 2022; Pennock et al., 2023; Jain et al., 2023). These prior reports of food-selective responses in the human brain have relied heavily on the same Natural Scenes Dataset (NSD) (Allen et al., 2022) of fMRI responses to a large set of natural images. Natural stimuli have the important advantage of relevance to real-world vision, but are also notoriously perilous due to the correlation in the natural world between high level concepts (like “face” or “food”) and low-level perceptual features (like “round” or “red”) (Norman-Haignere & McDermott, 2018; Harrison, 2021; Henderson et al., 2022). The prior studies reporting food selectivity put considerable effort into testing whether the neural response was selective for food per se, rather than its lower-level visual correlates. However, most of these control analyses were still performed on the same NSD data. Tests using more tightly controlled stimuli were limited, and in some cases were performed only on DNN-predicted responses rather than on actual fMRI measurements. Here, we moved beyond the space of NSD and DNN models to conduct stronger tests of the food selectivity hypothesis by measuring neural responses in new subjects to stimuli carefully constructed to unconfound the presence of food in the image from the visual features that tend to be present in natural images of food.

In both experiments, we first ran a functional localizer in each participant to identify the “food component” identified by Khosla et al (2022): a weighted set of voxels with strong responses to food images, which emerged from a data-driven analysis of the NSD data (see Methods and Figs. 1 and 2). We then measured the response of this food component to our new stimuli, collected across two experiments. In Experiment 1, because food is often colorful and appears less appetizing when color is removed (Lee et al., 2013; Ruseva et al., 2025), we tested greyscale versions of the food and non-food images from the NSD set. Second, to determine whether the food component response results from the food itself, rather than the background of NSD food images (e.g., table tops and plates), we created and measured responses to isolated food and non-food images on a blank white background, which we refer to as the “Cutout” condition. Third, because the food in the NSD images is generally within arm’s reach, often positioned in a table top scene, we tested whether our apparent food selectivity might derive in part from selectivity for Reachspaces (Josephs et al., 2021). In Experiment 2, we further tested the effect of distance, creating stimuli in which the same food is placed at two distances while holding the background constant. Next, because food often has soft material properties, we tested non-food items with a similar “gooey” texture. Further, because food items usually fall within a certain size range, we tested whether the food-selective response is driven by a preference for the real-world size of objects (Konkle & Oliva, 2012). Finally, we tested the role of mid-level perceptual features, like texture and curvature, using ‘Texform’ (Long et al., 2016) versions of the NSD food and non-food images, which preserve mid-level features while scrambling high-level semantic information.

**Fig. 1.**
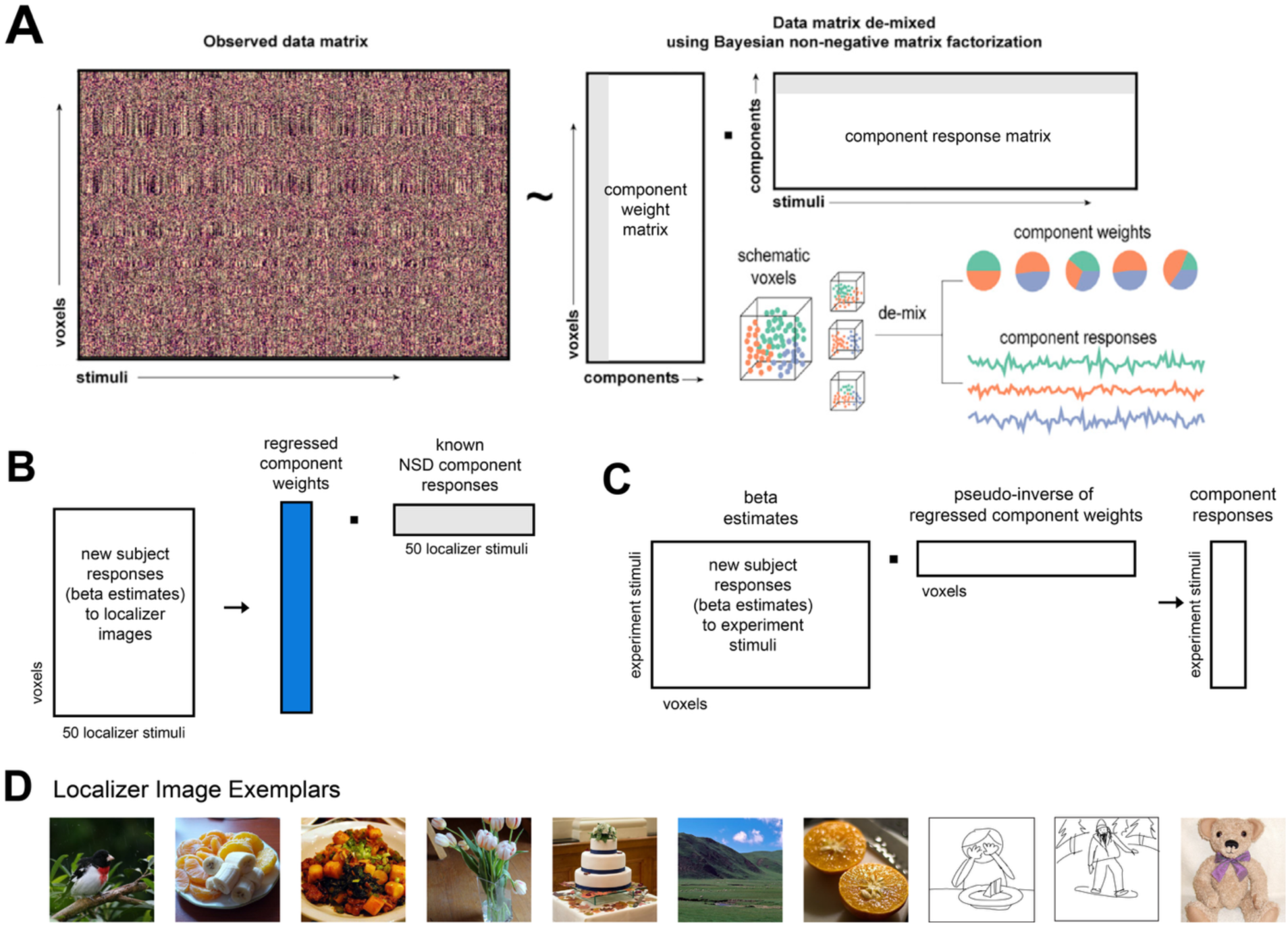
Food Component Analysis Pipeline. A) Schematic showing derivation of the food component from NSD subjects from Khosla et al (2022). The observed data matrix from NSD was factorized into a component weight matrix and a component response matrix. This approach “de-mixes” neural populations that cohabit voxels. Each component features a set of per-voxel weights (grey vertical bar in component weight matrix) as well as a component response to each stimulus image (grey horizontal bar in component response matrix). B) Deriving component weights (shown in blue) in new subjects. Using the new subject’s responses and the known NSD component responses to the same 50 localizer images, we use linear regression to derive the per-voxel component weights in each new subject (blue bar). C) Obtaining the component responses to new stimuli from our experiments. We multiply the matrix of per-voxel beta estimates for each experiment stimulus with the pseudoinverse of food component weights from the localizer (derived in part B). The resulting vector shows the inferred component response to each experiment stimulus. Parts B and C are done individually for each subject. D) Examples of localizer images. Photographs of people have been replaced with sketches.

**Fig. 2.**
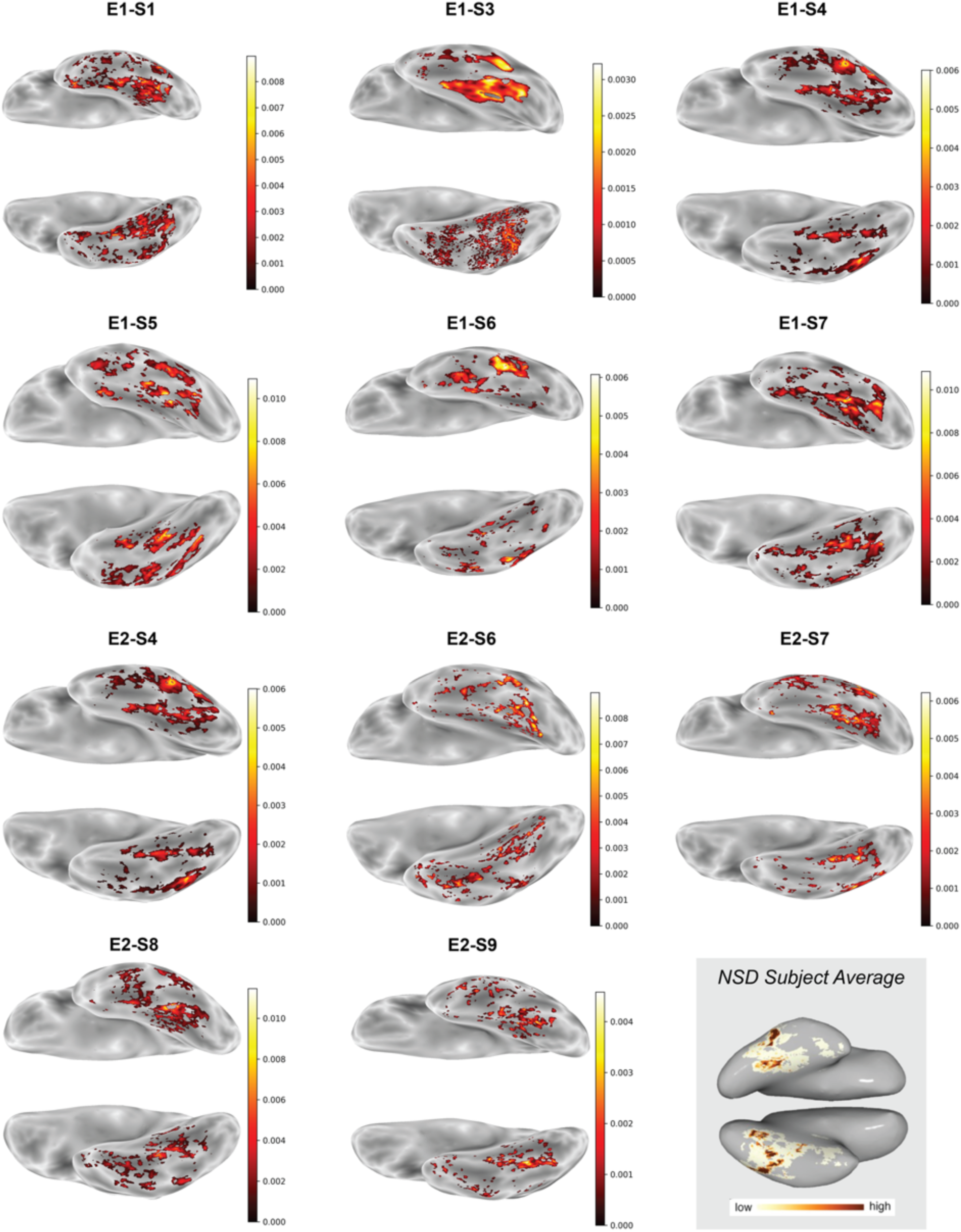
Component weight maps for all subjects from Experiments 1 and 2. Subjects 1-3 from Experiment 2 are not shown because they are identical to Experiment 1 subjects 1, 5, and 7 respectively. NSD subject average map from Khosla et al 2022 shown at bottom right.

In this work, we show that the “food component” is not strictly food selective, as this selectivity does not persist when the background is removed and visual features are matched, when the food is far away, and when the food is very small. Additionally, the component response to non-foods with a “gooey” material texture is just as high as its response to NSD food images. Our work highlights the necessity of using carefully controlled stimuli to test hypotheses derived from large naturalistic datasets, and suggests that the ventral visual pathway may contain highly replicable and robust response profiles that are not clearly interpretable.

## Results

### Experiment 1

We first successfully localized the food component in each participant, revealing the characteristic medial and lateral bands running along the posterior-anterior axis of the ventral pathway (Fig. 2), as reported in previous studies (Khosla et al., 2022; Pennock et al., 2023; Jain et al., 2023). We then measured the response of the food component to each of ten stimulus conditions, in a 2×5 design in which the food/non-food factor was crossed with five stimulus types. Stimulus types were i) original images from the NSD set, in color; ii) the same images converted to greyscale, iii) Cutout objects on a white background in color, iv) Cutout objects in greyscale, and v) Reachspace images (see Fig. 1 for examples). Within each stimulus type there were ten food images and ten non-food images. fMRI responses of the food component to each of the ten conditions, plus the fixation condition (averaging over the response to each of the ten images in each condition, and then across participants) are shown in Fig. 3B.

**Fig. 3.**
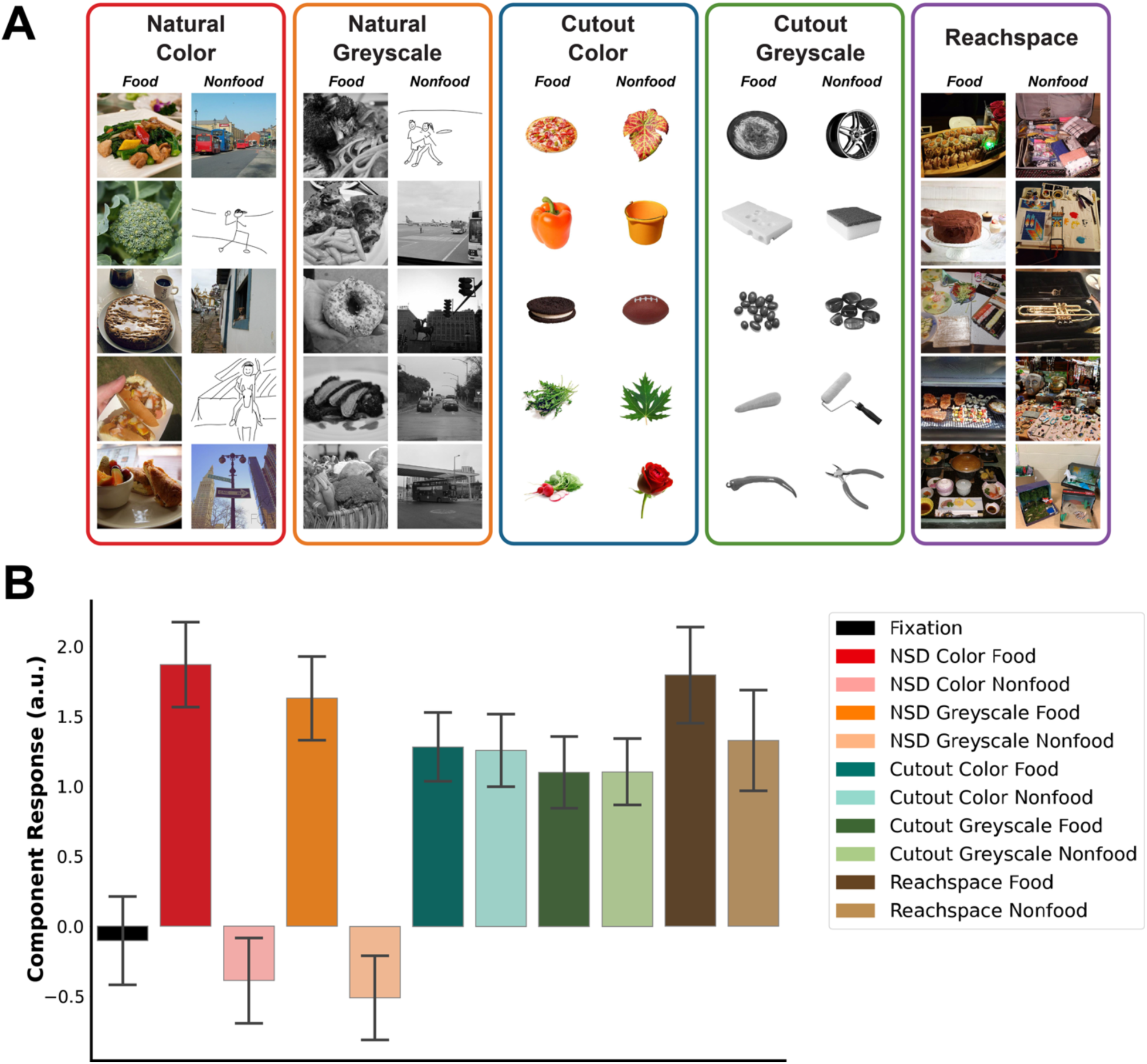
Conditions Tested and their Food Component Responses, Experiment 1. A) Example images from Experiment 1. Photographs of people have been replaced with sketches. B) Inferred magnitude of the food component response to each stimulus condition in Experiment 1, averaged across subjects (N=6). Note that responses are calculated on data independent from the food component localizer. Error bars show within-subject standard error.

Our first analysis model tested four pre-registered hypotheses about responses of the food component: 1) that we will replicate the originally reported (Khosla et al., 2022) higher response to food than non-food images from the NSD stimulus set, 2) that the response to color images will be higher than the response to greyscale images, 3) that there will be an interaction of food and color such that the food selectivity is greater when images are in color than in greyscale, and 4) that responses will be higher to food than non-food objects when both are isolated and pasted onto a white background (“Cutout”). To do so, we fit a mixed-effects model predicting activation to stimuli in all conditions except Reachspace and fixation. See Methods for more details about the models used in this section. Major findings are reported here with full results in Supp Table S1.

First, confirming Hypothesis 1, we successfully replicated the effect of Khosla et al. (2022), finding that the food component responded more strongly to NSD images of food than NSD images of non-food (estimate = 2.19, 95% CI = [1.89, 2.50], p <.001). For Hypothesis 2, we found a main effect of color across content and background, such that greyscale stimuli elicited significantly lower responses than color stimuli (estimate =-0.18, 95% CI = [-0.34, - 0.02], p =.033). As for Hypothesis 3, we found no significant interaction between food and color across background (estimate = 0.06, 95% CI = [−0.24, 0.37], p =.654). Lastly, for Hypothesis 4, we found no significant effect of food within the Cutout conditions, across both color and greyscale (estimate = 0.01, 95% CI = [−0.26, 0.29], p =.921), indicating that the effect of food is highly dependent on background. We then used Bayes Factors to investigate the statistical likelihood of the null hypothesis, i.e. that there is no effect of food within the Cutout condition. Across all priors, the Bayes Factors prefer a null effect of food within Cutout stimuli (Supp Table S3), ranging from 10.94 (strong evidence in favor of null, given a prior for a large effect) to 1.52 (anecdotal evidence in favor of null, given a prior for a very small effect). Thus, we replicated the effect of food within NSD images, found a higher response to color than greyscale images, no interaction between food and color, and no food selectivity when the food is removed from the background.

Next, we tested whether the food selectivity can be explained by a preference for Reachspaces, which predicts that we will find no difference between food and non-food if both are depicted in Reachspace images. Contrary to this hypothesis, we found a greater response to food over non-food within Reachspace images (estimate = 0.47, 95% CI = [0.08, 0.86], p =.018). However, we found a significant interaction between food and Reachspace such that the component prefers food over non-food more strongly in the NSD condition than the Reachspace condition (estimate = 1.77, 95% CI = [1.18, 2.36], p <.001).

To test whether our results are overly dependent on details of the component analysis, we also repeated these analyses using the average BOLD signal of the top 300 and 50 voxels with the highest food component weight in each subject, which is essentially a standard functional region of interest analysis. This analysis produced qualitatively similar findings, with full results in Supp Table S1. Finally, responses were analyzed separately within the top voxels of the medial and lateral bands, and showed qualitatively similar results, without notable differences between the two bands (Supp Fig. S4).

We also preregistered a number of additional analyses that aimed to better understand the functional and anatomical structure of this component. However, after finding that food selectivity did not persist when background was removed and was reduced when all stimuli were within reaching distance, we decided to focus our study instead on determining which features drive the large difference in component response between food and non-food in the NSD condition.

In sum, Experiment 1 replicated the original report of higher responses to food than non-food NSD images, and further found that this effect was not reduced in greyscale images, showing that the higher response to food than non-food images originally reported is not due to differences in the color content of food and non-food images. There was, however, a small but significantly higher response to color than greyscale NSD images, consistent with prior report (Pennock et al. 2023). Second, food preferences were small but significant when all stimuli depicted Reachspace scenes, showing that the original result cannot be entirely due to the fact that food was typically depicted in Reachspace scenes. Most importantly, however, we found no evidence of food selectivity when the background was removed (in the Cutout stimuli), challenging our prior claim that this component is primarily selective for food.

The lack of food selectivity for Cutout images was unexpected. The anatomical locations and functional response profile of this component are highly systematic, as reported in three prior studies. If this component is not selective for food, what is its function? We decided to step back and consider a broader space of hypotheses with a set of new exploratory analyses of the NSD data, our own behavioral experiments, and CNN-based models of the food component to test predictions of alternate hypotheses.

### Exploring alternative hypotheses with NSD data and computational models

The unexpected finding that food selectivity did not persist for Cutout images prompted us to seek alternative hypotheses that may explain the response profile of the component. We started by inspecting component responses to NSD images from the published data (Khosla et al. 2022) and identified a number of stimulus properties that appeared to be confounded with food in the NSD images (Fig. 4A) that might therefore explain or contribute to its response: the distance of the main objects and/or overall scene, the presence or affordance of hand actions (Jacobs et al., 2010; Bracci et al., 2012), real-world size (Konkle & Oliva, 2012), animacy (Konkle & Caramazza, 2013), and “gooey” material properties (Paulun et al., 2025). We explored these factors in two ways: First, we returned to the NSD shared 1000 images and collected human ratings on these dimensions. This enabled us to test i) which of these dimensions was correlated with the presence of food in NSD images, and ii) which dimensions might explain some or all of the response of the food component. Second, we used a predictive CNN model of the food component (from Khosla et al 2022) to test our alternative hypotheses with new stimuli. The analyses in this section were exploratory and not pre-registered.

**Fig. 4.**
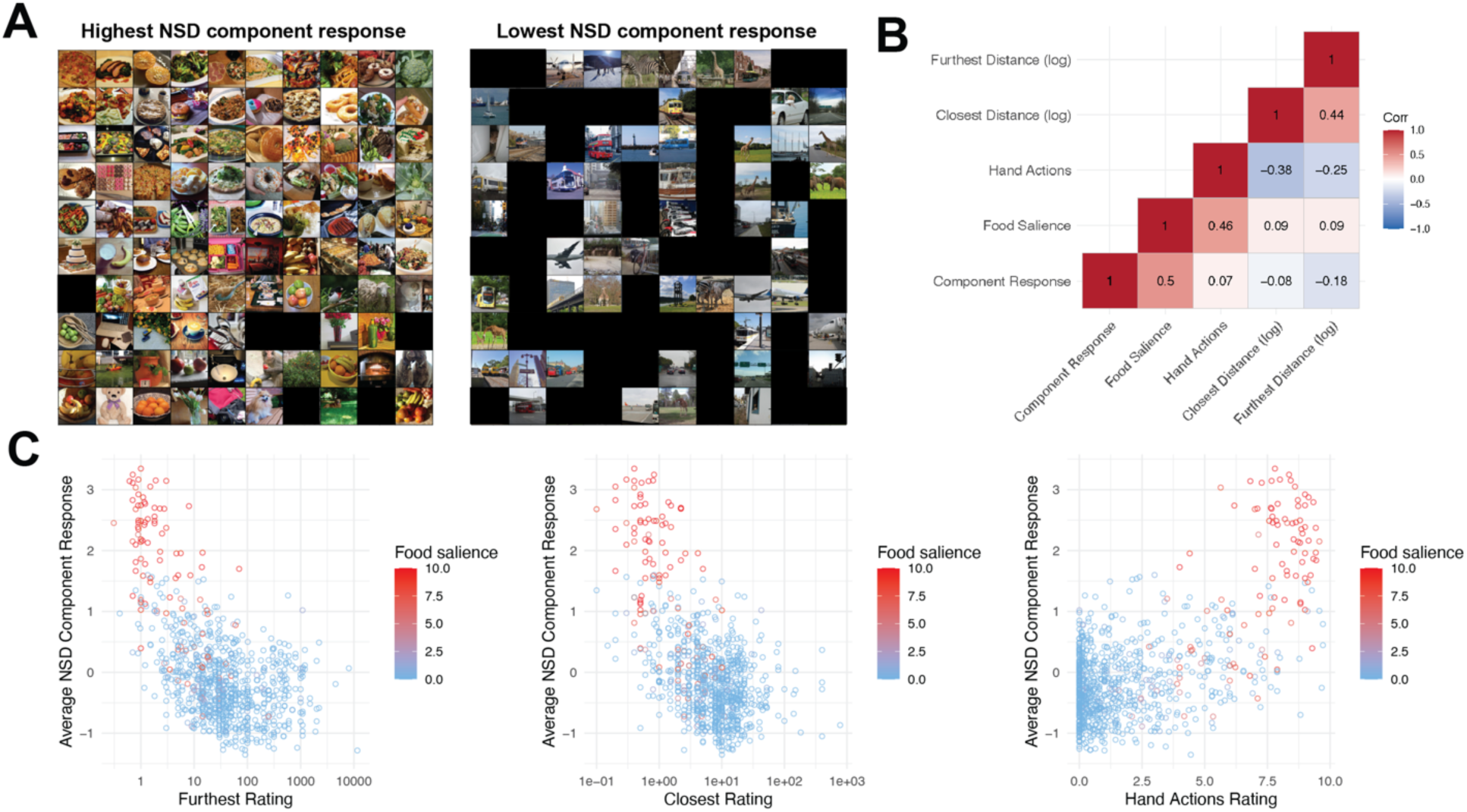
Exploring Alternative Possible Drivers of the Food Component Response. A) Top 100 and bottom 100 NSD images in food component response from Khosla et al (2022). Photographs of people have been redacted. B) Partial correlations of component response, food salience ratings, hand action salience ratings, log of closest distance ratings, and log of furthest distance ratings. All correlations are significant. C) Plots showing relationship between average NSD component response and (left to right) furthest distance ratings, closest distance ratings, and hand action salience ratings. Each point is one of the shared 1000 NSD images, and the color represents that image’s average food salience rating.

Our human ratings of food salience, foreground distance, background distance, and hand action affordance were all significantly correlated with one another across the 1000 shared NSD images (Fig. 4B), highlighting the difficulty of separating their effects in natural images.

Regression analyses showed that, despite independent contributions from both foreground and background distance, food salience consistently emerged as the strongest predictor of the food component response. We additionally found little evidence for an independent effect of hand action affordance. See Supp Section S6A for all results.

Next, to address the confounding image properties present in the NSD dataset, we used a highly predictive CNN model of the food component (correlation = 0.83 with held-out NSD data) (Khosla et al., 2022) to evaluate controlled, novel stimuli, serving as a way to “pilot” these stimuli prior to collecting fMRI data. We first tested the model on the stimuli from Experiment 1 and measured the correlation between model predictions and our real data, finding a correlation of r=0.63 with Experiment 1. While the model predicted a significant effect of food in the Cutout conditions, the effect size was much smaller than that of the NSD conditions. We then tested the model on novel stimuli, and it predicted that food no longer elicits a strong response when it is far from the camera, and that component responses decrease as background scenes become larger, but that this trend plateaus when scenes exceed the size of a large room. The model also preferred small over large objects and showed unexpectedly high responses to “gooey” non-food objects. See Supp Section S6B for all results. Overall, the above exploratory findings identified distance, real-world size, and gooeyness as candidate factors influencing the food component, motivating the controlled manipulations tested in Experiment 2.

### Experiment 2

Here we tested the most plausible alternative accounts of the selectivity of the food component that emerged from our exploratory analyses. To do this, we constructed a new set of controlled stimuli that unconfounded food from real-world size, distance, and gooeyness. We localized the food component in each subject following the same procedure as Experiment 1. We then scanned new participants (with the exception of 3 participants who also did Experiment 1) while they viewed the food component localizer images, and then the new set of stimuli, and measured the response of the food component. The stimulus design and analysis methods were preregistered before running the full experiment.

Responses of the food component to each of the 13 conditions are shown in Fig. 5B. The mixed-effects analysis models used for Experiment 2 mirror those used in Experiment 1, with details in Methods. Major findings are reported here with all results in Supp Table S2. First, we once again replicated the significant effect of food within NSD images (estimate = 2.71, p<.001, 95% CI [1.83, 3.58]). We then tested whether mid-level visual features might explain the component response (Long et al., 2018, Kramer et al., 2023), and found that the response to Texform Food is not significantly different from Texform Non-food (estimate = 0.28, p = 0.349, 95% CI [-0.31, 0.88]). Importantly, we did not find conclusive evidence of a difference between NSD Food and Texform Food (estimate = 0.37, p =.279, 95% CI [-0.30, 1.04],), suggesting that the recognizability of food may not be necessary for the high food response. Bayes Factors show anecdotal evidence in favor of the null (see SI for Bayes Factors on Top 300 and 50 voxels, which show anecdotal evidence against the null). Conversely, a further exploratory analysis showed that the response to NSD Non-food was significantly lower than Texform Non-food (estimate=-2.05, p<.001, 95% CI [-3.04,-1.06]), indicating that recognizability or some aspect of the structure of the natural non-food images is surprisingly necessary for the low non-food response. Thus, the original contrast of NSD food and NSD non-food is likely driven in part by unusually low responses to natural non-food.

**Fig. 5.**
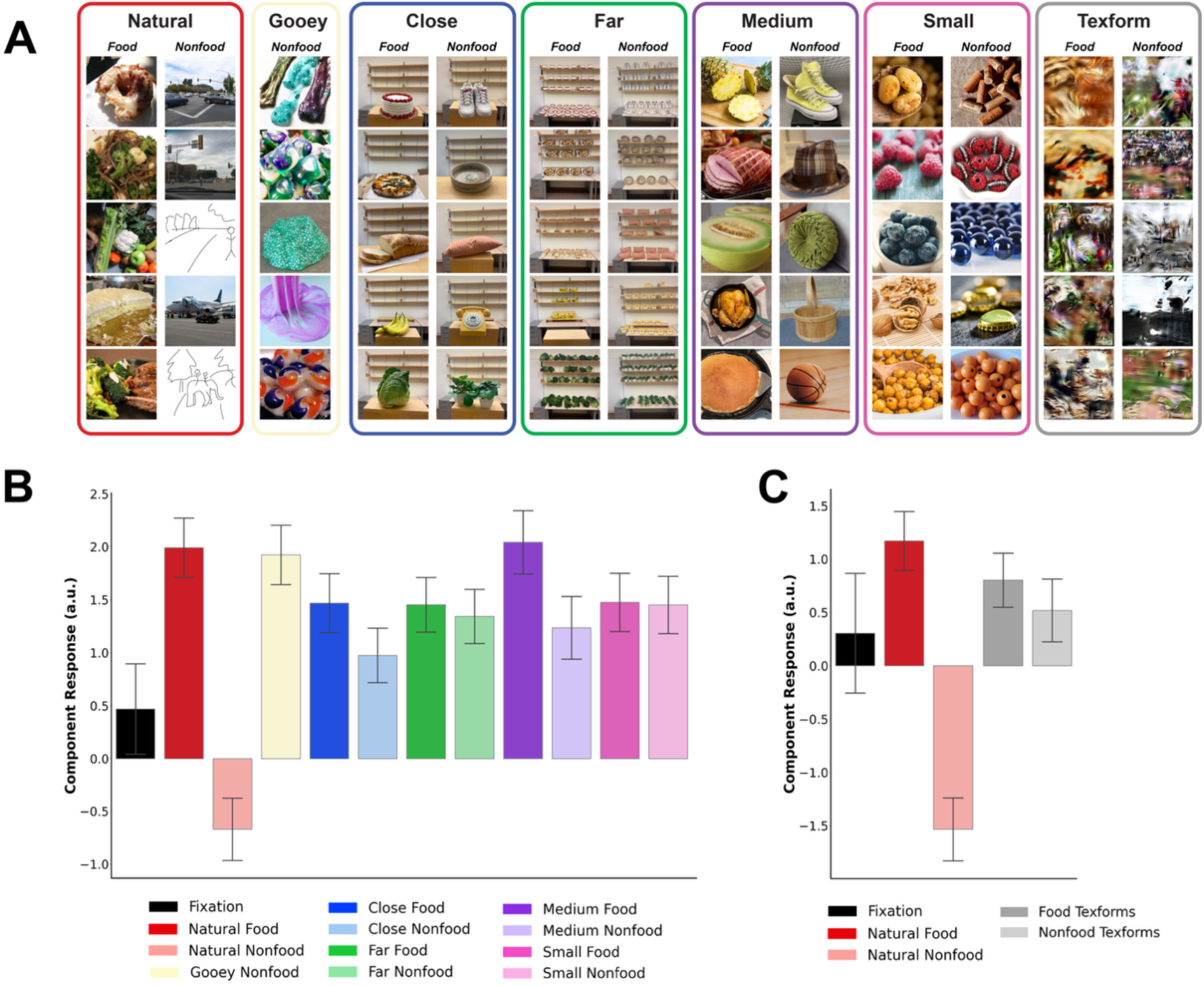
Stimulus conditions and Results for Experiment 2. A) Examples of stimuli from Experiment 2. Photographs of people have been replaced with sketches. B) Inferred magnitude of component response to each stimulus condition in Experiment 2, averaged across subjects (N=8). Error bars show within-subject standard error. The Texform condition is not included because only 6 subjects saw the Texform condition. C) Component response to Texform conditions, averaged over subjects who saw the Texform condition (N=6). See Supp Fig. S4B for plot with Texforms and all other conditions.

Next, we tested the effect of the distance of the focal object. The exploratory analyses in the previous section showed that distance was a predictor of component response independent of food salience; additionally, CNN model analyses predicted that the food component would not have a selective response to food when it was far away from the camera. On average, across both Close and Far images, food elicited a higher activation than non-food (estimate = 0.30, p = 0.018, 95% CI [0.06, 0.55]), whereas distance, across both food and non-food, did not have a significant effect (estimate =-0.18, p = 0.174, 95% CI [-0.44, 0.08]). Although we did not find a significant interaction between food and distance (estimate = 0.38, p = 0.124, 95% CI [-0.11, 0.88]), the effect of food was larger within Close images relative to Far images. Bayes Factors supported a null effect of food within Far images, with a range of values between 7.14 and 1.16, depending on prior. Thus, the food component is not selective for a given distance overall, but food selectivity is absent for distant food.

Next, we tested the effects of real-world size, and found on average, between Small and Medium images, a higher response to food than non-food (estimate = 0.42, p =.027, 95% CI [0.05, 0.78]) and a significant interaction such that there was a smaller effect of food for Small than Medium (estimate =-0.78, p =.007, 95% CI [-1.34,-0.23]). We did not find a significant main effect of real-world size across food and non-food (estimate =-0.18, p =.281, 95% CI [-0.51, 0.15]). We found a higher response to food than non-food in the Medium condition (estimate = 0.81, p<.001, 95% CI [0.37, 1.24]) but not in the Small condition (estimate = 0.02, p =.912, 95% CI [-0.41, 0.46]). Bayes Factors supported a null effect of food within Small images, with a range of values between 6.46 and 1.19 depending on prior. Thus, we find no evidence of an independent effect of real-world size, and additionally find that food selectivity does not persist when the food is very small.

Lastly, we tested the effect of material gooeyness, and found that the component response to Small Food was significantly lower than that of Gooey Non-food (estimate =-0.45, p =.028, 95% CI [-0.84,-0.05]), and that there was no significant difference between NSD Food and Gooey Non-food (estimate = 0.07, p =.747, 95% CI [-0.37, 0.51]). Bayes Factors supported the null effect of NSD Food vs. Gooey Non-food, ranging from 6.35 to 1.20 based on priors. This analysis was not preregistered as we only preregistered a comparison between Gooey Non-food and Small Food.

We repeated all the above analyses using the average BOLD signal over the Top 300 and 50 voxels with the highest component weight for each subject, as well as in medial and lateral bands separately, and found qualitatively similar results (Supp Table S2, Fig S5).These analyses were preregistered except for those indicated as exploratory analyses or specified otherwise.

Altogether, the results of Experiment 2 support the finding from Experiment 1 that the food component is not selective for food alone, as it also has a similarly high response to Gooey Non-food and to behaviorally unrecognizable Texform images. Additionally, the distance of the focal object does not explain component response, but food selectivity goes away when the food or non-food is far away from the camera. We find significant food selectivity in two planned contrasts (Close food versus non-food and Medium food versus non-food), but the effect size for these comparisons is far smaller than for the original NSD food and non-food images. A greater challenge to the food selectivity hypothesis comes from the lack of food selectivity for the Far or Small conditions, and the fact that the response to Gooey Non-food and Texform Food were not significantly different than the response to NSD food. Thus, although some conditions show a degree of selectivity for food, many do not, and it is clear that this component is not selectively responsive to food alone.

Our findings challenge the notion that this component is selective for the semantic category of food. Instead, it might be selective for a certain set of image statistics that tend to be present in food images. To gain some insight into the low-level features or image statistics that differentiate food from other categories, we passed the 1000 shared NSD images through AlexNet, and then performed PCA on responses of layer units, and found that food images clustered in AlexNet fc6, with emergence of these groups in the conv2 layer. These analyses were exploratory and not preregistered. Additional details and plots are in Supp Section S7.

We hypothesize that if there are visual statistics that are common in food, and that the “food component” is responsive to these visual statistics, it might function as a first-level filter to detect whether the visual input is likely to be food, before sending the information to a later processing stage that might more definitively detect or analyze food. If this were the case, then we would expect it to take longer to recognize non-food as non-food if it has a high food component response, and conversely it would take longer to recognize food as food if it has a low food component response. We tested this using an online behavioral experiment in which participants were shown each of the 1000 shared NSD images, and they were asked to make a judgment of whether it was food or non-food as fast as possible. Consistent with our hypothesis, we found a negative correlation between RT and component response in food images (r = - 0.48, p < 0.001) and a positive correlation between RT and component response in non-food images (r = 0.38, p < 0.001). We also collected RTs for our controlled stimulus sets from Experiments 1 and 2, but did not see clear trends of correlation between RT and component response (Supp Fig. S8). Together, these results suggest that the food component response captures visual features that facilitate food categorization in naturalistic images, but that this relationship between RT and component response does not generalize to out-of-distribution images in which many of these visual features are reduced or removed.

Inspired by our finding from Experiment 1 that there was no effect of food in the Cutout condition, we conducted an additional exploratory analysis of the 1000 shared NSD images in order to investigate whether the component response could be explained by just the background, rather than the food itself. We selected images from NSD that did not contain food, but that featured either “food contexts” such as kitchens and tabletops, or “non-food contexts” of similarly sized indoor scenes, approximately matched for visual features. We did not find a significant difference between food contexts and non-food contexts in NSD data (Supp Fig. S9).

## Discussion

Despite reports of food selectivity in visual cortex from 3 papers, and extensive efforts to rule out alternative accounts of the selectivity of the “food component” (Khosla et al. 2022), we show here that the “food component” fails many tests of classic category selectivity. Specifically, although we replicate in both of our experiments the strong food selectivity found with the NSD stimuli used in previous reports, we find much weaker albeit significant food selectivity in several new conditions here (Reachspace Food, Close Food, and Medium-sized Food), and no food selectivity when the background is removed, when the food is far from the camera, or when the real-world size of the food is very small. Even more problematic for the food selectivity hypothesis is the finding that “food component” responses to Gooey Non-food and to unrecognizable Texforms were as high as to NSD food. Thus, the “food component” is not selective solely for the semantic category of food, but is both context-and material-sensitive, and we will refer to it henceforth as the “food-ish component”. Our findings were qualitatively similar when analyzing the food-ish component and when measuring responses of the most food-selective voxels, and similar for the medial and lateral bands (see Supp Fig. S5). Revisiting the data from NSD led us to consider distance as a potential predictor, as the NSD images with the highest component response are predominantly close to the camera and the images with the lowest component response are predominantly large outdoor scenes. However, both our regression analyses of NSD data and our results from Experiment 2 show that neither foreground nor background distance can explain the response of the food-ish component. We thus tested many alternative hypotheses of what this component might be selective for, but did not find a single interpretable account of the response profile. Beyond rejecting the hypothesis that the ventral pathway contains a neural population selective for only food, this study highlights the risk of drawing conclusions based on only natural stimuli, and it raises the possibility that some highly replicable and robust response profiles in the ventral visual pathway may simply not be straightforwardly interpretable with a single clear semantic label (like faces or scenes or bodies).

We have focused on the methods and results from Khosla et al. (2022), but food selectivity has previously been reported by two other groups. Jain et al. (2023) reported significant food selective voxels in new subjects using a contrast of food > faces, places, bodies, and words using Cutout greyscale images of these categories presented on a scrambled background. They computed the voxel-wise selectivity (computed as the difference between the response to the preferred category divided by the average response to the non-preferred categories) of food-selective voxels and found that similar cortical regions responded more to food than non-food in this contrast as in the contrast from the NSD data. These findings are consistent with ours in finding higher responses to food than non-food even for greyscale images, but contrast with our finding of no food selectivity for Cutout images. This discrepancy could reflect either the presence of a scrambled background in Jain’s study, or the fact that their non-food images were not closely matched to the food images for visual features (as they were for the Cutout images in our study).

Another study by Pennock et al. (2023) identified color-biased voxels within NSD subjects and found that food images were the strongest predictor of activity in these regions. Their finding is consistent with our result from Experiment 1 in which we found a significant effect in the component response to color. Pennock’s color-biased regions were identified by finding voxels whose responses were significantly correlated with average color saturation; we conducted a similar analysis of our data and found a small but significant correlation between saturation and component response (Experiment 1: r = 0.16, p = 0.038; Experiment 2: r = 0.38, p < 0.001). Given that we still found an effect of food within the greyscale NSD condition, color is not crucial to the response profile of this component, which is consistent with Pennock’s finding that the presence of food in the image was the strongest predictor of activity in their food-preferring voxels.

Another recent study, Ritchie et al. (2024) has suggested that food selectivity may be explained by selectivity for tools, after finding significant overlap between voxels that preferred tools over animals and voxels that preferred food over animals. In our exploratory analyses of NSD data, we found that the salience of hand actions in an image (which is related to tool-ness, in that tools are likely to have a very high hand action rating) was a much weaker predictor of component response than food salience. Across three different model fittings that used ratings of food salience, salience of hand actions, and foreground and background distance as predictors of component response, hand actions had a consistently smaller beta value than food salience, including one model in which hand actions were not a significant predictor.

Additionally, the Ritchie et al study focuses on graspable foods, while the food images most preferred by the component in our study are often close-ups of non-graspable foods such as pasta and rice. Thus, the food preferences observed in our study are not attributable to hand actions or tools.

Our work highlights the methodological principle that because naturalistic stimuli contain inherent confounds between high-level categories and low-level visual features, findings that use large-scale datasets of naturalistic images must be complemented with carefully controlled experiments that isolate the features of interest. Our additional analyses of the NSD images showed a high correlation of NSD responses to distance and food salience. Despite extensive control analyses in Khosla et al. (2022), all empirical measurements in that study were from the same NSD images in which features were confounded. It was only with carefully selected stimuli controlling for specific features that we were able to fully unconfound food salience from these other dimensions.

Even though the food-ish component is not selective for only food, it is robust and replicable, with an anatomically consistent location across participants. How then might we characterize it? Across all our data, the most consistent finding was a very low response to NSD non-food images, which was true both for the component response and the top voxel response. The NSD non-food images consist primarily of outdoor scenes, frequently featuring human bodies, animals, or large vehicles. Across both experiments, all conditions were within the same ballpark response magnitude as the NSD food images, except for natural non-food which was much lower. This can also be seen in the Cutout and Texform conditions, in which Cutout food and Texform food are both just as high as NSD food, but Cutout nonfood and Texform nonfood are much higher than NSD nonfood. Removing the background or scrambling the image does not reduce the response to food, but rather to increases the response to non-food. Thus, we find a neural population in the brain whose response profile is robust and replicable, and yet so far resists simple interpretation, and is largely defined by what it does *not* like, rather than what it likes (Gondur et al., 2026). Our work also suggests that dimensions of representation in the ventral pathway may not be easily interpretable, reflecting visual image statistics that are highly correlated with, but not congruent to, semantic categories in the real world, consistent with past findings that relatively primitive visual features such as curvature can explain some of the large-scale topographical organization of the ventral pathway (Long et al., 2018). Lastly, we restate the importance of designing experiments with carefully controlled stimuli in order to test hypotheses. Despite the many advantages of hypothesis-free data-driven approaches with large datasets of naturalistic stimuli, it is impossible to unconfound many factors in these naturalistic images, and thus they should not by themselves be used to draw robust conclusions.

## Limitations

Our study has two main limitations. First, while our localizer method is able to infer the food-ish component in new subjects, these component weights are not as accurate as if subjects saw all the NSD images and then voxel decomposition was used independently in each subject to derive the top components. Additionally, the component weights in new subjects are estimated using linear regression on natural images, and thus might not be robust to out-of-distribution or “non-natural” images such as our Cutout stimuli. Our component-localization approach also assumes a degree of consistency in the component response profile across individuals, such that component responses estimated in one group of participants (the NSD subjects) can be used to infer component weights in new participants. Prior work (Khosla et al., 2022) has shown that while food-ish component responses are broadly consistent across individuals, they are not perfectly correlated and exhibit inter-subject variability. As a result, our method may underestimate individual differences in food selectivity or obscure idiosyncratic response patterns. However, scanning each participant on the full set of images used in NSD would require vastly more time, and we think our method strikes a good balance between time and accuracy. Our method enables us to robustly replicate the previously reported much higher response to NSD food than NSD non-food, suggesting that the shorter component localizer is effective.

Second, many of our conditions which were not food selective (such as Cutout and Far) had a small difference in means in the correct direction, and may reach significance if many more subjects were scanned, despite Bayes Factors supporting a null effect. However, even if these conditions produced significant food selectivity with a much larger number of participants, it is already clear that the magnitude of food selectivity in those conditions is much lower than for NSD images.

### General Methods

The stimulus design and analysis methods were preregistered before running the full experiment.

Experiment 1: [https://osf.io/w4nq3].

Experiment 2: [https://osf.io/rcyq4].

### Participants

In Experiment 1, seven participants (ages 22 to 30, 4 female) participated in the fMRI experiment, each for three two-hour sessions. We initially preregistered six subjects based on a power analysis of pilot data. However, one subject reported red/green colorblindness after participating in the experiment, and their data were excluded from analysis, and we thus scanned a seventh participant.

In Experiment 2, eight participants (ages 22 to 30, 5 female) participated in the fMRI experiment. Three of the participants for Experiment 2 also participated in Experiment 1. N=6 subjects were preregistered. However, after we ran the first two subjects, there was an update to the experiment, in which we changed two conditions from pixel scrambled food/non-food to Texform food/non-food, and thus there were N=6 subjects who completed the second version of Experiment 2.

Before participating in the experiment, all participants gave informed consent to the experimental protocol approved by the MIT Committee on the Use of Humans as Experimental Subjects (no. 0403000096). The study was conducted in compliance with all the relevant ethical guidelines and regulations for work with human participants.

### Procedure

Both Experiments 1 and 2 consisted of three 2-hour scan sessions for each participant. The first session consisted of a high-resolution anatomical scan and 20 runs of the food localizer, which consisted of fifty images, each shown a total of 20 times. Each image was presented in an event-related design for 300ms and followed by minimum inter-stimulus-interval (ISI) of 3700ms and maximum ISI of 11700ms optimized using OptSeq2 (https://surfer.nmr.mgh.harvard.edu/optseq/). There was no overt task; subjects were simply instructed to fixate on a small 0.3dva fixation cross at the center of the screen.

In both Experiments 1 and 2, the second and third session each consisted of 10 runs each of the main experiment for a total of 20 runs. In Experiment 1, each run consisted of 100 stimuli presented once each, along with an additional fixation condition. In Experiment 2, each run consisted of 104 stimuli presented once each, along with an additional fixation condition. In both experiments, each image was shown a total of 20 times across the whole experiment.

Both experiments were run in an event-related design, with the stimulus conditions (10 in Experiment 1, 13 in Experiment 2), plus a fixation condition, presented in a different pseudorandomized order (using OptSeq) within each run. Each image was presented for 300ms and followed by minimum inter-stimulus-interval (ISI) of 3700ms and maximum ISI of 11700ms optimized using OptSeq2. Subjects performed a one-back task during the experiment, such that one image was selected randomly per run within each condition to be repeated for a one-back trial. These were analyzed as “targets” and not as instances of the stimulus categories they were derived from. Throughout the experiment, subjects were instructed to fixate on a small fixation cross 0.3 degrees of visual angle at the center of the screen.

In both Experiments 1 and 2, each participant completed three 2-hour scan sessions, with the following exceptions: in Experiment 1, one subject completed Experiment 2 first, and thus we used that participant’s localizer data from Experiment 2. In Experiment 2, two subjects also participated in Experiment 1 and thus did not repeat the food localizer and only completed the latter two 2-hour scan sessions.

### Experiment 1 Stimuli

Experiment 1 tested the effect of color, background, and Reachspace on the food component response across ten stimulus conditions in a 2×5 design in which the food/non-food factor was crossed with five stimulus types (See Fig. 3A), creating ten stimulus conditions. Stimulus types were i) original images from the NSD set, in color; ii) the same images converted to greyscale, iii) Cutout objects on a white background in color, iv) Cutout objects in greyscale, and v) Reachspace images (see Fig. 1 for examples). Within each stimulus type there were ten food images and ten non-food images. All stimuli used in the study were pre-registered.

The NSD food and non-food images were selected from the Natural Scenes Dataset, such that the food images had a high food component response in NSD data in our earlier study and the non-food images had a low food component response (Khosla et al., 2022). The non-food images largely consisted of outdoor scenes, often with a person, animal, or large object in the foreground. The NSD greyscale food and non-food conditions were the same images, converted to greyscale. To eliminate participants’ ability to mentally “fill in” the color from their memory of the color version of the greyscale images, the images were counterbalanced across subjects such that half the subjects saw set A in color and set B in greyscale, and the other half of subjects saw the opposite. The Cutout images were images of food and non-food from the FoodPics dataset (Blechert et al., 2014), with pictures of food and non-food isolated on a white background. Food/non-food pairs were matched for visual features by selecting pairs that had maximum cosine similarity between vector representations in the penultimate layer of a CNN. This was done using both AlexNet (Krizhevsky et al., 2012) and ResNet (He et al., 2016), and then of these visually matched pairs, we hand selected the stimuli for the study to maximize semantic and feature variability between images within each category. As with the natural images, we used two sets of Cutout stimuli so that half the participants saw set A in color and set B in greyscale, and half saw the opposite. The Reachspace images were taken from the Reachspace Database (Josephs et al., 2021), which feature images of items and surfaces within arms’ reach, like tabletops, desks, and sinks. From this database we selected 10 Reachspace scenes clearly containing food and 10 clearly not containing food. These images were selected in matched pairs to control for low-level visual features, such that food and non-food scenes produced similar representations in a pretrained AlexNet.

### Experiment 2 Stimuli

Experiment 2 tested the effect of mid-level features, distance, real-world size, and material gooeyness on the food component response across thirteen stimulus conditions in a 2 x 6 + 1 design, in which the food/non-food factor was crossed with image type (see Fig. 5a), which had 6 levels: NSD images, Texform scrambled versions of these NSD images, Close and Far (to manipulate Distance), Small and Medium in Real-world size; an additional single Gooey Non-food condition was included. There were eight images in each condition for a total of 104 images.

First, to replicate the food selectivity we originally reported with NSD images, and to provide benchmarks against which to measure responses in the other conditions, we included the natural NSD food and non-food conditions in color. Because we had some overlapping subjects between Experiment 1 and Experiment 2, we chose images from NSD that had high similarity to the NSD food and non-food images from Experiment 1 when comparing image representations in the penultimate layer of Resnet. To test whether mid-level visual features present in food might explain the response even in a stimulus that was not recognizable as food, we made scrambled “Texform” versions of the NSD food and non-food images from Experiment 1, generated according to Long et al. (2016). We confirmed in an online behavioral test that food could not be distinguished from non-food in the Texform versions of these images.

The Close and Far conditions tested the effect of distance of the focal object by maintaining the same background while changing the distance of the focal object. While both foreground and background distance were significant predictors of component response in NSD data, here we chose to vary the distance of the food/non-food rather than the distance of the background because our CNN model predicted no effect of food when it is placed a mid-distance away from the camera, and also predicted that changing background distance by altering the size of a scene would not have a large effect on the component response. These Close and Far stimuli were photographed in-house and featured food and non-food objects, approximately matched in shape, size, and color. The Close condition showed a single food or non-food object a few inches from the camera, and the Far condition showed the same object in the same background setting but 6-8 feet away from the camera. To approximately control for pixel count, the Close condition depicted one or few objects and the Far condition depicted many instances of the same object. As a result of this design, the two conditions differ in numerosity, and the Far condition involves clutter and repetition not featured in the Close condition, a compromise that was necessary in order to control for pixel count. Participants verbally confirmed they were able to identify the pictured food and non-food items in the images. To test the effect of real-world size, we found images online that featured items around the size of a coin or raspberry (the Small condition), or items around the size of a melon or shoe (the Medium condition). The food and non-food objects in these conditions were approximately balanced for color and shape, and objects were approximately balanced between the small and medium conditions for pixel count. Finally, to test whether the food component was sensitive to the material property of gooeyness, we included a Gooey Non-food condition containing closeup images of colorful slime, Tide pods, etc., which our CNN model predicted a high response to.

This condition was designed to be contrasted with the visually similar small non-food condition.

### Component Localizer Design

A 50-stimulus food component localizer was developed to localize the food component in the new subjects. Deriving components de novo requires very large amounts of data, since both the component response profiles and the voxel weights must be jointly inferred, typically relying on statistics computed across thousands of images and multiple participants. Instead, for the localizer scan, we leveraged the food component identified in our earlier work (Khosla et al., 2022) and measured fMRI responses in new participants to a subset of 50 NSD images for which the component responses were already known. To make the localizer efficient, we selected the 50 NSD images using the stimulus-selection approach from Boebinger et al. (2026). Briefly, starting from the full NSD image set, we greedily removed images while preserving the subset that minimized the expected variance of the inferred component weights. This procedure favors images that provide complementary information about the known component response profiles, allowing participant-specific voxel weights to be estimated from a much smaller localizer set. Exemplars can be seen in Fig. 1d.

### fMRI data acquisition

All imaging was performed on a Siemens 3T Prisma scanner with a 32-channel head coil at the Athinoula A. Martinos Imaging Center at the Massachusetts Institute of Technology (MIT). For each participant, a high-resolution T1-weighted anatomical image [Magnetization Prepared RApid Gradient-Echo (MPRAGE): repetition time (TR) = 2.53 s, echo time (TE) = 3.57 ms, α = 9°, field of view (FOV) = 256 mm, matrix = 256 × 256, slice thickness = 1 mm, 176 slices, acceleration factor = 2, 24 reference lines, bandwidth (BW) = 190 Hz per pixel] was collected in addition to whole-brain functional data using a T2*-weighted echo planar imaging pulse sequence (TR = 2 s, TE = 30 ms, α = 90°, FOV = 1836 mm, matrix = 918 × 918, slice thickness = 2 mm, voxel size = 2 mm by 2 mm in plane, slice gap = 0 mm, 66 slices).

### fMRI data preprocessing

Preprocessing was done using FreeSurfer (https://freesurfer.net/). FMRI data preprocessing included motion correction, slice time correction, linear fit to detrend the time series, and spatial smoothing with a Gaussian kernel (FWHM = 2 mm). Before smoothing the functional data, all functional runs were co-registered to the subject’s T1-weighted anatomical image. A GLM was run on each participant individually using GLMsingle (Prince et al., 2022) including the experimental conditions (10 conditions in Experiment 1 and 13 conditions in Experiment 2) and 6 nuisance regressors based on the motion estimates (x, y, and z translation; roll, pitch, and yaw of rotation). Analyses were performed on the subject’s native inflated cortical surface (using FreeSurfer’s mri_vol2surf function).

### Inferring Component Response to New Experimental Conditions

The food voxel weights in new participants were inferred from the known NSD component responses to each image using linear regression (Fig. 1b). The food voxel weights were then used to measure the response of the inferred food component to the conditions tested in each experiment (Fig. 1c). Formally, to estimate each participant’s component weights, we multiplied that participant’s data matrix of voxel responses to the 50 selected natural image stimuli (***D*** ∈ ℝ^images×voxels^) by the pseudoinverse of the known component response matrix for those same images (***R*** ∈ ℝ^images×components^), yielding voxel-wise component weights *W*^ = ***R***⁺***D***. These weights were then used as predictors in a second linear regression: for each experimental stimulus, we estimated the component response amplitudes that best reconstructed the observed voxel-response pattern across cortex (***R***_test_ = ***D***_test_ *W*^⁺). This procedure is analogous to identifying voxels responsive to a localizer contrast and subsequently measuring their responses to a separate set of experimental stimuli and has been used in prior work for localizing music-selective components (Boebinger et al., 2026). The localizer was run independently from the main experiment. The inferred component responses of the localizer had a Fisher z-averaged correlation of 0.78 with the average component responses of NSD subjects.

### Analysis Models

Once the food component response was estimated for each stimulus in each participant as explained above, we then conducted statistical tests of our hypotheses. All of the analyses were preregistered with the exception of those explicitly stated as exploratory. However, we did not preregister the specific type of model (linear mixed-effects model), the contrast coding, or packages used, and determined this later. The analysis models used are Gaussian mixed-effects linear models using the R (R Core Team, 2024) packages lme4 (Bates et al., 2015, version 1.1.37) and lmerTest (Kuznetsova et al., 2017, version 3.1.3). All models used a maximal random effect structure for the grouping variables of participants and image (Barr, 2013). This means that we used random intercepts for participants and images, with all the fixed effects as random slopes within participants and a random slope for greyscale within image where relevant. All reported models have converged, meaning that the results are stable and reliable. After fitting each mixed-effects model, we also computed model-estimated marginal means using the R emmeans package (Lenth & Piaskowski, 2026, version 1.11.2.8). Degrees of freedom for model coefficients were computed using the Satterthwaite approximation (lmerTest); estimated marginal means and their contrasts use asymptotic degrees of freedom, as all models exceeded the default sample-size limit for exact methods in emmeans. All reported contrasts compare two conditions and are presented without correction for multiple comparisons. To compute Bayes Factors, we used Savage-Dickey method (Wagenmakers et al., 2010) on fits from the brms package (Bürkner, 2017, version 2.22.0), which had a comparable fixed and random effects structure to the maximum likelihood fits from lme4 unless stated otherwise. To use comparable priors across experiments, we z-scored the response variable within each analysis, so that all effects are expressed in units of the standard deviation of the response in that analysis. This was necessary because the component response and the top-voxel responses differ substantially in scale (SD ≈ 2.7–3.0 and ≈ 0.4–0.6, respectively), so a prior specified in raw units would correspond to a very different effect size in each. Since Bayes Factors are dependent on priors (Schad et al., 2023), we selected a range of 3 meaningful priors for the predictors that we tested for a null effect: each prior was Normal(0, sd), where sd was 0.5, 0.2, and 0.05, placing ∼95% of the prior mass within ±1.0 (large), ±0.4 (medium), and ±0.1 (negligible) SD of the response, respectively. The priors on all other parameters were held fixed across the three models and were weakly informative: Normal(0, 1) on all remaining population-level coefficients, Student-t(3, 0, 2.5) on the intercept, on the random-effect standard deviations, and on the residual standard deviation, and an LKJ(1) prior on the random-effect correlation matrices. All models were fit with 4 chains of 8,000 iterations (2,000 warmup), yielding 24,000 post-warmup samples, with adapt_delta = 0.95 and a fixed seed, unless stated otherwise in the model-specific sections below. All models converged, with R̂ ≤ 1.002 and bulk ESS > 2,900. Details for each model are described below.

### Experiment 1 Models

Analysis of Food/Non-food × Color/Greyscale × NSD/Cutout on the Eight Stimulus Conditions Generated by Orthogonally Crossing these Factors: Mixed-effects model with three sum-coded binary factors: food/non-food, color/greyscale, and Cutout/NSD (food=-0.5, non-food =0.5; color=-0.5, greyscale=0.5; Cutout=0.5, NSD=-0.5). Because the color and greyscale versions of an image are the same photograph, they share an image identifier, making greyscale the only factor that varies within an image. The subsequent pairwise comparisons of estimated marginal means compared one pair of conditions within another pair of conditions, averaged over the third pair. For instance, we compared food to non-food separately within NSD images and within Cutout images, collapsing across color and greyscale. The Bayesian models used the same settings but with the data filtered to Cutout only, thus dropping the background predictor from the model.

Analysis of Food/Non-food and NSD/Reachspace on the Four Stimulus Conditions Generated by Orthogonally Crossing these Factors: Mixed-effects model with two treatment-coded binary factors: food/non-food, and NSD/Reachspace (food=0, non-food=1; NSD=0, Reachspace=1). We used treatment coding because the Reachspace manipulation affected food and non-food images in opposite directions (food images were made more distant, whereas non-food images were made closer), so the average main effects of food/non-food or NSD/Reachspace would not have a meaningful interpretation.

### Experiment 2 Models

Analysis of Food/Non-food and NSD/Texform on the Four Stimulus Conditions Generated by Orthogonally Crossing these Factors: Mixed-effects model with two treatment-coded binary factors: food/non-food, and NSD/Texform (food=0, non-food=1; natural=0, scrambled=1).

Similar to the Food/Reachspace model described above, we chose treatment coding because our primary question was how Texform scrambling altered responses within food and within non-food images, rather than whether there was an overall mean effect of scrambling across image categories. The Bayesian models used the same data and settings; the effect tested was the Texform coefficient, which under this coding is the difference between Texform Food and NSD Food.

Analysis of Food/Non-food and Close/Far (Distance) on the Four Stimulus Conditions Generated by Orthogonally Crossing these Factors: Mixed-effects model with two sum-coded binary factors: food/non-food and close/far (food=0.5, non-food=-0.5; close=0.5, far=-0.5). The Bayesian models used the same settings but with the data filtered to Far only, thus dropping the distance predictor from the model; This model used adapt_delta = 0.99.

Analysis of Food/Non-food and Small/Medium (Real-world Size) on the Four Stimulus Conditions Generated by Orthogonally Crossing these Factors: Mixed-effects model with two sum-coded binary factors: food/non-food and small/medium (food = 0.5, non-food =-0.5; medium =-0.5, small = 0.5). The Bayesian models used the same settings but with the data filtered to Small only, thus dropping the real-world size predictor from the model; This model used adapt_delta = 0.99.

Analysis of Gooey Non-food, NSD Food, and Small Food: Mixed-effects model with one 3-level factor of condition, with Gooey non-food as the reference level. We did not perform a pairwise comparison of estimated marginal means for this model. This model is not preregistered as we preregistered a contrast between Gooey Non-food and Small Food only. The Bayesian models used the same data and settings; the effect tested was the NSD Food coefficient, which under this coding is the difference between NSD Food and Gooey Non-food.

## Acknowledgments.

This work was supported by McGovern Institute Seed Funds.

## Declaration of Interests

The authors declare no competing interests.

## Notes

### Competing Interest Statement

The authors have declared no competing interest.

## References

Allen, E. J., St-Yves, G., Wu, Y., Breedlove, J. L., Prince, J. S., Dowdle, L. T., Nau, M., Caron, B., Pestilli, F., Charest, I., et al. (2022). A massive 7T fMRI dataset to bridge cognitive neuroscience and artificial intelligence. Nature Neuroscience, 25(1), 116--126.

Barr, D. J. (2013). Random effects structure for testing interactions in linear mixed-effects models. In Frontiers in psychology (Vol. 4, p. 328). Frontiers Media SA.

Bates, D., Mächler, M., Bolker, B., & Walker, S. (2015). Fitting linear mixed-effects models using LME4. Journal of Statistical Software, 67(1), 1--48. 10.18637/jss.v067.i01

Blechert, J., Meule, A., Busch, N. A., & Ohla, K. (2014). Food-PICS: An image database for experimental research on eating and appetite. Frontiers in Psychology, 5, 617.

Boebinger, D., McDermott, J. H., Kanwisher, N., & Norman-Haignere, S. (2026). Music-selective cortex is sensitive to structure in both pitch and time. Cerebral Cortex, 36(6), bhag072.

Bracci, S., Cavina-Pratesi, C., Ietswaart, M., Caramazza, A., & Peelen, M. V. (2012). Closely overlapping responses to tools and hands in left lateral occipitotemporal cortex. Journal of Neurophysiology, 107(5), 1443--1456.

Bürkner, P.-C. (2017). BRMS: An R package for bayesian multilevel models using Stan. Journal of Statistical Software, 80, 1--28. 10.18637/jss.v080.i01

Gondur, R., Stan, P., Smith, M. A., & Cowley, B. (2026). A tale of two tails: Preferred and anti-preferred natural stimuli in visual cortex. International Conference on Learning Representations, 2026, 98177--98204.

Harrison, W. J. (2021). Luminance and contrast of images in the THINGS database. Perception, 51, 244–262. https://api.semanticscholar.org/CorpusID:235789357

He, K., Zhang, X., Ren, S., & Sun, J. (2016). Deep residual learning for image recognition. Proceedings of the IEEE Conference on Computer Vision and Pattern Recognition, 770--778.

Henderson, M. M., Tarr, M. J., & Wehbe, L. (2022). Low-level tuning biases in higher visual cortex reflect the semantic informativeness of visual features. Journal of Vision, 23. https://api.semanticscholar.org/CorpusID:251474277

Jacobs, S., Danielmeier, C., & Frey, S. H. (2010). Human anterior intraparietal and ventral premotor cortices support representations of grasping with the hand or a novel tool. Journal of Cognitive Neuroscience, 22(11), 2594--2608.

Jain, N., Wang, A., Henderson, M. M., Lin, R., Prince, J. S., Tarr, M. J., & Wehbe, L. (2023). Selectivity for food in human ventral visual cortex. Communications Biology, 6(1), 175.

Josephs, E. L., Zhao, H., & Konkle, T. (2021). The world within reach: An image database of reach-relevant environments. Journal of Vision, 21(7), 14--14.

Kanwisher, N. (2025). Animal models of the human brain: Successes, limitations, and alternatives. Current Opinion in Neurobiology, 90, 102969.

Khosla, M., Murty, N. A. R., & Kanwisher, N. (2022). A highly selective response to food in human visual cortex revealed by hypothesis-free voxel decomposition. Current Biology, 32(19), 4159--4171.

Konkle, T., & Caramazza, A. (2013). Tripartite organization of the ventral stream by animacy and object size. The Journal of Neuroscience, 33(25), 10235--10242.

Konkle, T., & Oliva, A. (2012). A real-world size organization of object responses in occipitotemporal cortex. Neuron, 74(6), 1114--1124.

Kramer, L. E., Chen, Y.-C., Long, B., Konkle, T., & Cohen, M. R. (2023). Contributions of early and mid-level visual cortex to high-level object categorization. bioRxiv.

Krizhevsky, A., Sutskever, I., & Hinton, G. E. (2012). Imagenet classification with deep convolutional neural networks. Advances in Neural Information Processing Systems, 25.

Kuznetsova, A., Brockhoff, P. B., & Christensen, R. H. B. (2017). lmerTest package: Tests in linear mixed effects models. Journal of Statistical Software, 82(13), 1--26. 10.18637/jss.v082.i13

Lee, S.-M., Lee, K.-T., Lee, S.-H., & Song, J.-K. (2013). Origin of human colour preference for food. Journal of Food Engineering, 119(3), 508--515.

Lenth, R. V., & Piaskowski, J. (2026). Emmeans: Estimated marginal means, aka least-squares means. https://rvlenth.github.io/emmeans/

Long, B., Konkle, T., Cohen, M. A., & Alvarez, G. A. (2016). Mid-level perceptual features distinguish objects of different real-world sizes. Journal of Experimental Psychology: General, 145(1), 95.

Long, B., Yu, C.-P., & Konkle, T. (2018). Mid-level visual features underlie the high-level categorical organization of the ventral stream. Proceedings of the National Academy of Sciences, 115(38), E9015--E9024.

Norman-Haignere, S. V., & McDermott, J. H. (2018). Neural responses to natural and model-matched stimuli reveal distinct computations in primary and nonprimary auditory cortex. PLoS Biology, 16(12), e2005127.

Paulun, V. C., Pramod, R. R. T., Tenenbaum, J. B., & Kanwisher, N. (2025). Dissociable cortical regions represent things and stuff in the human brain. Current Biology, 35(17), 4075-- 4083.

Pennock, I. M., Racey, C., Allen, E. J., Wu, Y., Naselaris, T., Kay, K. N., Franklin, A., & Bosten, J. M. (2023). Color-biased regions in the ventral visual pathway are food selective. Current Biology, 33(1), 134--146.

Prince, J. S., Charest, I., Kurzawski, J. W., Pyles, J. A., Tarr, M. J., & Kay, K. N. (2022). Improving the accuracy of single-trial fMRI response estimates using GLMsingle. Elife, 11, e77599.

R Core Team. (2024). R: A language and environment for statistical computing. R Foundation for Statistical Computing. https://www.R-project.org/

Ritchie, J. B., Andrews, S. T., Vaziri-Pashkam, M., & Baker, C. I. (2024). Graspable foods and tools elicit similar responses in visual cortex. Cerebral Cortex, 34(9), bhae383.

Ruseva, D., Giesel, M., & Hesse, C. (2025). Fading appetite: Desaturation of food images reduces cravings but not approach biases. Appetite, 108323.

Schad, D. J., Nicenboim, B., Bürkner, P.-C., Betancourt, M., & Vasishth, S. (2023). Workflow techniques for the robust use of bayes factors. Psychological Methods, 28(6), 1404.

Wagenmakers, E.-J., Lodewyckx, T., Kuriyal, H., & Grasman, R. (2010). Bayesian hypothesis testing for psychologists: A tutorial on the savage--dickey method. Cognitive Psychology, 60(3), 158--189.

