## Supplemental for "Apparent food selectivity reflects multiple non-food image properties"

### Supplemental Table S1. Results of Analyses of Responses of Food Component in Experiment 1

#### a1. Analysis of Food/Non-food × Color/Greyscale × NSD/Cutout on the Eight Stimulus Conditions Generated by Orthogonally Crossing these Factors: Fixed Effects

| | Component Response | | | | Top 300 Voxels | | | | Top 50 Voxels | | | |
|---|---|---|---|---|---|---|---|---|---|---|---|---|
| Term | Estimate | SE | p-value | CI | Estimate | SE | p-value | CI | Estimate | SE | p-value | CI |
| Food | -1.10 | 0.10 | < .001 | [-1.31, -0.90] | -0.14 | 0.02 | < .001 | [-0.18, -0.10] | -0.19 | 0.03 | < .001 | [-0.24, -0.13] |
| Color | -0.18 | 0.07 | 0.033 | [-0.34, -0.02] | -0.03 | 0.02 | 0.162 | [-0.07, 0.01] | -0.05 | 0.03 | 0.183 | [-0.12, 0.03] |
| Cutout | 0.54 | 0.24 | 0.067 | [-0.05, 1.12] | 0.03 | 0.03 | 0.425 | [-0.05, 0.10] | 0.02 | 0.05 | 0.643 | [-0.10, 0.14] |
| Food x Color | 0.06 | 0.13 | 0.654 | [-0.24, 0.37] | -0.01 | 0.02 | 0.583 | [-0.07, 0.04] | -0.02 | 0.03 | 0.476 | [-0.10, 0.05] |
| Food x Cutout | 2.18 | 0.22 | < .001 | [1.72, 2.64] | 0.28 | 0.04 | < .001 | [0.19, 0.37] | 0.37 | 0.06 | < .001 | [0.23, 0.50] |
| Color x Cutout | 0.03 | 0.11 | 0.815 | [-0.20, 0.25] | -0.00 | 0.02 | 0.973 | [-0.04, 0.04] | 0.02 | 0.03 | 0.420 | [-0.04, 0.09] |
| Food x Color x Cutout | -0.06 | 0.25 | 0.827 | [-0.61, 0.50] | -0.08 | 0.04 | 0.087 | [-0.17, 0.01] | -0.11 | 0.05 | 0.080 | [-0.24, 0.02] |

#### a2. Pairwise Contrasts of Food versus Non-food, Color versus Greyscale, and NSD versus Cutout

| | | Component Response | | | | Top 300 Voxels | | | | Top 50 Voxels | | | |
|---|---|---|---|---|---|---|---|---|---|---|---|---|---|
| Condition | Contrast | Estimate | SE | p-value | CI | Estimate | SE | p-value | CI | Estimate | SE | p-value | CI |
| Cutout | Food - Non-food | 0.01 | 0.14 | 0.921 | [-0.26, 0.29] | -0.00 | 0.03 | 0.983 | [-0.05, 0.05] | 0.00 | 0.03 | 0.910 | [-0.06, 0.07] |
| NSD | Food - Non-food | 2.19 | 0.16 | < .001 | [1.89, 2.50] | 0.28 | 0.03 | < .001 | [0.23, 0.34] | 0.37 | 0.04 | < .001 | [0.29, 0.45] |
| Food | Cutout - NSD | -0.55 | 0.26 | 0.036 | [-1.07, -0.04] | -0.11 | 0.04 | 0.004 | [-0.19, -0.04] | -0.16 | 0.06 | 0.010 | [-0.28, -0.04] |
| Non-food | Cutout - NSD | 1.63 | 0.27 | < .001 | [1.10, 2.15] | 0.17 | 0.03 | < .001 | [0.10, 0.24] | 0.21 | 0.05 | < .001 | [0.11, 0.30] |
| Food | Color - Greyscale | 0.21 | 0.10 | 0.029 | [0.02, 0.40] | 0.02 | 0.02 | 0.338 | [-0.02, 0.06] | 0.03 | 0.04 | 0.342 | [-0.04, 0.10] |
| Non-food | Color - Greyscale | 0.15 | 0.09 | 0.107 | [-0.03, 0.33] | 0.03 | 0.02 | 0.063 | [-0.00, 0.07] | 0.06 | 0.03 | 0.062 | [-0.00, 0.12] |

#### b1. Analysis of Food/Non-food and NSD/Reachspace on the Four Stimulus Conditions Generated by Orthogonally Crossing these Factors: Fixed Effects

| | Component Response | | | | Top 300 Voxels | | | | Top 50 Voxels | | | |
|---|---|---|---|---|---|---|---|---|---|---|---|---|
| Term | Estimat e | SE | p-value | CI | Estimate | SE | p-value | CI | Estimate | SE | p-value | CI |
| Food | -1.35 | 0.12 | < .001 | [-1.60, -1.11] | -0.20 | 0.03 | < .001 | [-0.26, -0.13] | -0.26 | 0.04 | < .001 | [-0.35, -0.16] |
| Reachspace | 0.82 | 0.26 | 0.017 | [0.20, 1.44] | 0.08 | 0.03 | 0.050 | [0.00, 0.16] | 0.08 | 0.04 | 0.104 | [-0.02, 0.18] |
| Food x Reachspace | 1.77 | 0.28 | < .001 | [1.18, 2.36] | 0.20 | 0.04 | 0.001 | [0.11, 0.29] | 0.26 | 0.06 | 0.004 | [0.12, 0.40] |

### Supplemental Table S1 (cont.)

### b2. Pairwise Contrasts of Food versus Non-food and NSD versus Reachspace

| | | Component Response | | | | Top 300 Voxels | | | | Top 50 Voxels | | | |
|---|---|---|---|---|---|---|---|---|---|---|---|---|---|
| Condition | Contrast | Estimate | SE | p-value | CI | Estimate | SE | p-value | CI | Estimate | SE | p-value | CI |
| NSD | Food - Non-food | 2.24 | 0.17 | < .001 | [1.91, 2.57] | 0.30 | 0.03 | < .001 | [0.23, 0.36] | 0.39 | 0.05 | < .001 | [0.29, 0.48] |
| Reachspace | Food - Non-food | 0.47 | 0.20 | 0.018 | [0.08, 0.86] | 0.10 | 0.03 | 0.005 | [0.03, 0.16] | 0.13 | 0.05 | 0.008 | [0.03, 0.23] |
| Food | NSD - Reachspace | 0.07 | 0.26 | 0.798 | [-0.45, 0.58] | 0.02 | 0.02 | 0.423 | [-0.03, 0.07] | 0.05 | 0.03 | 0.088 | [-0.01, 0.11] |
| Non-food | NSD - Reachspace | -1.70 | 0.32 | < .001 | [-2.34, -1.07] | -0.18 | 0.05 | < .001 | [-0.27, -0.09] | -0.21 | 0.06 | 0.001 | [-0.33, -0.08] |

### Supplemental Table S2. Results of Analyses of Responses of Food Component in Experiment 2

#### a1. Analysis of Food/Non-food and NSD/Texform on the Four Stimulus Conditions Generated by Orthogonally Crossing these Factors: Fixed Effects

| | Component Response | | | | Top 300 Voxels | | | | Top 50 Voxels | | | |
|---|---|---|---|---|---|---|---|---|---|---|---|---|
| Term | Estimate | SE | p-value | CI | Estimate | SE | p-value | CI | Estimate | SE | p-value | CI |
| Food | -2.71 | 0.45 | < .001 | [-3.68, -1.73] | -0.36 | 0.06 | 0.001 | [-0.52, -0.21] | -0.50 | 0.07 | < .001 | [-0.67, -0.32] |
| Texform | -0.37 | 0.34 | 0.292 | [-1.08, 0.34] | -0.11 | 0.05 | 0.042 | [-0.22, -0.01] | -0.17 | 0.06 | 0.030 | [-0.32, -0.02] |
| Food × Texform | 2.42 | 0.56 | < .001 | [1.22, 3.62] | 0.35 | 0.07 | < .001 | [0.20, 0.50] | 0.48 | 0.07 | < .001 | [0.31, 0.65] |

#### a2. Pairwise Contrasts of Food versus Non-food and NSD versus Texform

| | | Component Response | | | | Top 300 Voxels | | | | Top 50 Voxels | | | |
|---|---|---|---|---|---|---|---|---|---|---|---|---|---|
| Condition | Contrast | Estimate | SE | p-value | CI | Estimate | SE | p-value | CI | Estimate | SE | p-value | CI |
| NSD | Food - Non-food | 2.71 | 0.45 | < .001 | [1.83, 3.58] | 0.36 | 0.06 | < .001 | [0.24, 0.49] | 0.50 | 0.07 | < .001 | [0.36, 0.64] |
| Texform | Food - Non-food | 0.28 | 0.30 | 0.349 | [-0.31, 0.88] | 0.01 | 0.03 | 0.654 | [-0.04, 0.06] | 0.02 | 0.04 | 0.604 | [-0.05, 0.09] |
| Food | NSD - Texform | 0.37 | 0.34 | 0.279 | [-0.30, 1.04] | 0.11 | 0.05 | 0.014 | [0.02, 0.20] | 0.17 | 0.06 | 0.007 | [0.05, 0.29] |
| Non-food | NSD - Texform | -2.05 | 0.51 | < .001 | [-3.04, -1.06] | -0.24 | 0.05 | < .001 | [-0.34, -0.14] | -0.31 | 0.05 | < .001 | [-0.40, -0.21] |

#### b1. Analysis of Food/Non-food and Close/Far (Distance) on the Four Stimulus Conditions Generated by Orthogonally Crossing these Factors: Fixed Effects

| | Component Response | | | | Top 300 Voxels | | | | Top 50 Voxels | | | |
|---|---|---|---|---|---|---|---|---|---|---|---|---|
| Term | Estimate | SE | p-value | CI | Estimate | SE | p-value | CI | Estimate | SE | p-value | CI |
| Food | 0.30 | 0.12 | 0.018 | [0.06, 0.55] | 0.05 | 0.02 | 0.025 | [0.01, 0.09] | 0.06 | 0.03 | 0.020 | [0.01, 0.12] |
| Distance | -0.18 | 0.13 | 0.174 | [-0.44, 0.08] | -0.04 | 0.02 | 0.180 | [-0.09, 0.02] | -0.04 | 0.03 | 0.312 | [-0.11, 0.04] |
| Food x Distance | 0.38 | 0.24 | 0.124 | [-0.11, 0.88] | 0.07 | 0.04 | 0.117 | [-0.02, 0.16] | 0.09 | 0.06 | 0.155 | [-0.04, 0.22] |

#### b2. Pairwise Contrasts of Food versus Non-food and Close versus Far

| | | Component Response | | | | Top 300 Voxels | | | | Top 50 Voxels | | | |
|---|---|---|---|---|---|---|---|---|---|---|---|---|---|
| Condition | Contrast | Estimate | SE | p-value | CI | Estimate | SE | p-value | CI | Estimate | SE | p-value | CI |
| Close | Food - Non-food | 0.49 | 0.18 | 0.005 | [0.15, 0.84] | 0.08 | 0.03 | 0.011 | [0.02, 0.15] | 0.11 | 0.04 | 0.013 | [0.02, 0.20] |
| Far | Food - Non-food | 0.11 | 0.16 | 0.500 | [-0.21, 0.43] | 0.01 | 0.03 | 0.650 | [-0.04, 0.06] | 0.02 | 0.04 | 0.582 | [-0.05, 0.09] |
| Food | Close - Far | 0.02 | 0.18 | 0.933 | [-0.34, 0.37] | 0.00 | 0.04 | 0.980 | [-0.08, 0.08] | 0.01 | 0.05 | 0.855 | [-0.10, 0.12] |
| Non-food | Close - Far | -0.37 | 0.16 | 0.026 | [-0.69, -0.04] | -0.07 | 0.03 | 0.006 | [-0.12, -0.02] | -0.08 | 0.03 | 0.019 | [-0.15, -0.01] |

### Supplemental Table S2 (cont.)

#### c1. Analysis of Food/Non-food and Small/Medium (Real-world size) on the Four Stimulus Conditions Generated by Orthogonally Crossing these Factors: Fixed Effects

| | Component Response | | | | Top 300 Voxels | | | | Top 50 Voxels | | | |
|---|---|---|---|---|---|---|---|---|---|---|---|---|
| **Term** | **Estimate** | **SE** | **p-value** | **CI** | **Estimate** | **SE** | **p-value** | **CI** | **Estimate** | **SE** | **p-value** | **CI** |
| Food | 0.42 | 0.17 | 0.027 | [0.05, 0.78] | 0.07 | 0.04 | 0.137 | [-0.02, 0.16] | 0.09 | 0.06 | 0.202 | [-0.05, 0.22] |
| RW Size | -0.18 | 0.16 | 0.281 | [-0.51, 0.15] | -0.01 | 0.03 | 0.708 | [-0.06, 0.04] | -0.00 | 0.03 | 0.960 | [-0.07, 0.07] |
| Food x RW Size | -0.78 | 0.27 | 0.007 | [-1.34, -0.23] | -0.10 | 0.05 | 0.045 | [-0.19, -0.00] | -0.12 | 0.06 | 0.049 | [-0.24, -0.00] |

#### c2. Pairwise Contrasts of Food versus Non-food and Small versus Medium

| | | Component Response | | | | Top 300 Voxels | | | | Top 50 Voxels | | | |
|---|---|---|---|---|---|---|---|---|---|---|---|---|---|
| **Condition** | **Contrast** | **Estimate** | **SE** | **p-value** | **CI** | **Estimate** | **SE** | **p-value** | **CI** | **Estimate** | **SE** | **p-value** | **CI** |
| Medium | Food - Non-food | 0.81 | 0.22 | < .001 | [0.37, 1.24] | 0.11 | 0.05 | 0.031 | [0.01, 0.22] | 0.14 | 0.07 | 0.053 | [-0.00, 0.29] |
| Small | Food - Non-food | 0.02 | 0.22 | 0.912 | [-0.41, 0.46] | 0.02 | 0.04 | 0.669 | [-0.06, 0.10] | 0.03 | 0.06 | 0.672 | [-0.09, 0.15] |
| Food | Medium - Small | 0.57 | 0.21 | 0.007 | [0.15, 0.98] | 0.06 | 0.04 | 0.099 | [-0.01, 0.13] | 0.06 | 0.04 | 0.171 | [-0.03, 0.15] |
| Non-food | Medium - Small | -0.22 | 0.21 | 0.299 | [-0.62, 0.19] | -0.04 | 0.03 | 0.244 | [-0.10, 0.03] | -0.06 | 0.04 | 0.185 | [-0.14, 0.03] |

#### d1. Analysis of Gooey Non-food, NSD Food, and Small Food: Fixed Effects*

| | Component Response | | | | Top 300 Voxels | | | | Top 50 Voxels | | | |
|---|---|---|---|---|---|---|---|---|---|---|---|---|
| **Term** | **Estimate** | **SE** | **p-value** | **CI** | **Estimate** | **SE** | **p-value** | **CI** | **Estimate** | **SE** | **p-value** | **CI** |
| Small Food | -0.45 | 0.19 | 0.028 | [-0.84, -0.05] | -0.04 | 0.03 | 0.258 | [-0.10, 0.03] | -0.05 | 0.05 | 0.265 | [-0.15, 0.04] |
| NSD Food | 0.07 | 0.21 | 0.747 | [-0.37, 0.51] | 0.02 | 0.04 | 0.686 | [-0.06, 0.10] | 0.02 | 0.05 | 0.732 | [-0.09, 0.13] |

*This is a three-level factor, treatment coded with Gooey Non-food as the reference level. Each beta estimate indicates the effect of that condition in reference to Gooey Non-food.

### Supplemental Table S3. Bayes Factors for Effects Tested Against a Null

| | Component Response | | | Top 300 Voxels | | | Top 50 Voxels | | |
|---|---|---|---|---|---|---|---|---|---|
| **Effect Size Prior (Z-scored)** | **0.5** | **0.2** | **0.05** | **0.5** | **0.2** | **0.05** | **0.5** | **0.2** | **0.05** |
| Cutout Food – Cutout Non-food | 10.94 | 4.58 | 1.52 | 8.32 | 3.54 | 1.27 | 8.56 | 3.62 | 1.35 |
| NSD Food – Texform Food | 2.52 | 1.34 | 1.01 | 0.72 | 0.53 | 0.89 | 0.41 | 0.40 | 0.85 |
| Far Food – Far Non-food | 7.14 | 2.87 | 1.16 | 6.58 | 2.76 | 1.21 | 6.93 | 3.04 | 1.24 |
| Small Food – Small Non-food | 6.46 | 2.75 | 1.19 | 3.96 | 1.94 | 1.07 | 3.70 | 1.70 | 1.07 |
| NSD Food – Gooey Non-food | 6.35 | 2.75 | 1.20 | 4.50 | 1.97 | 1.07 | 4.51 | 1.96 | 1.10 |

Values are BF01, so values above 1 indicate evidence in favor of the null (no effect) and values below 1 indicate evidence in favor of a difference. Column headings give the standard deviation of the Normal(0, SD) prior placed on the effect tested; the response was z-scored within each analysis, so priors and effects are in units of the standard deviation of the response.

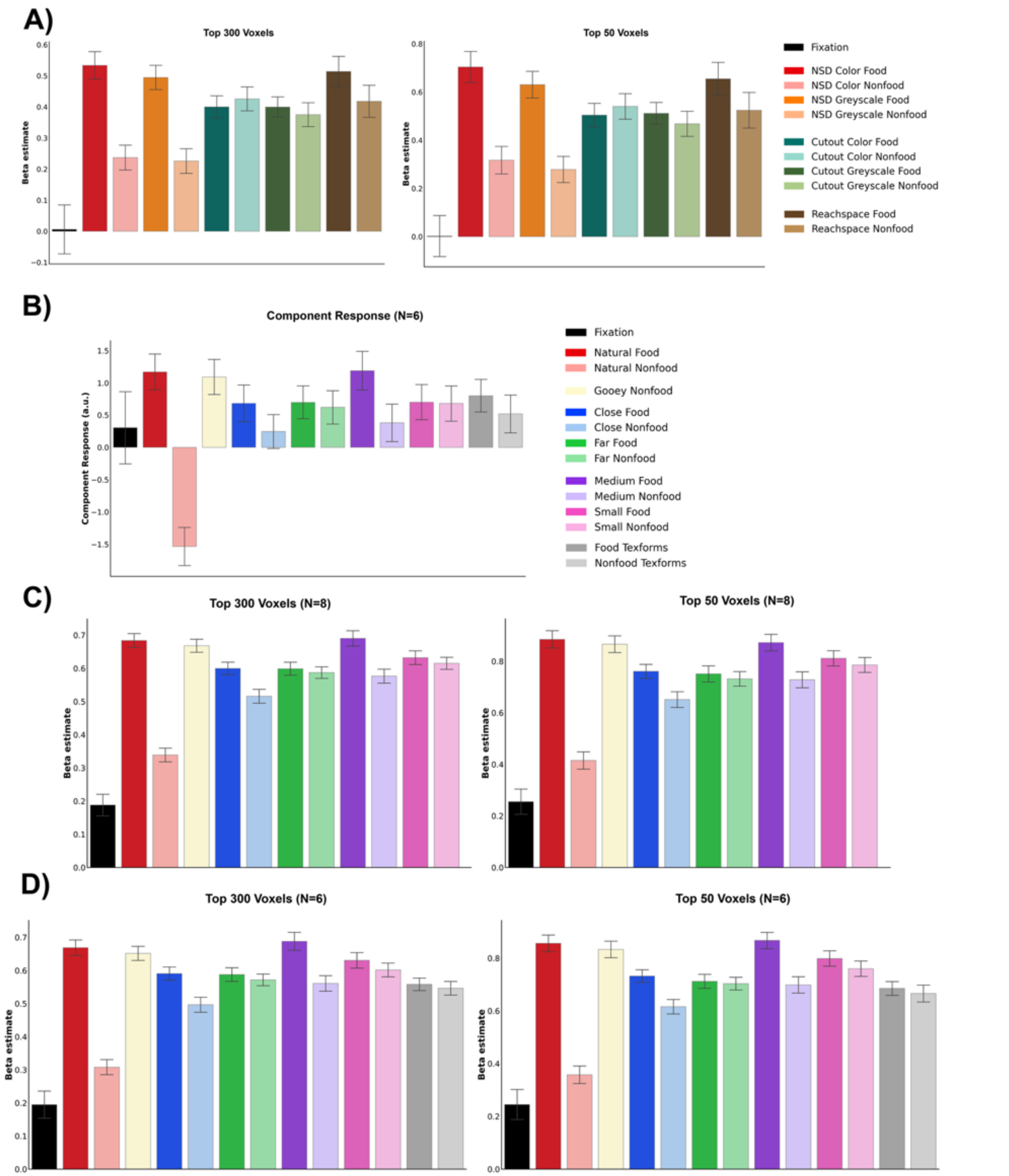


**Supplemental Fig. S4.** (A) Beta responses to conditions from Experiment 1, averaged over 300 (left) or 50 (right) voxels with the highest component weights. (B) Component response to all conditions from Experiment 2, including the Texform conditions. Data from only 6 subjects is included in this plot because only 6 subjects saw the Texform conditions. (C) Beta responses to conditions from Experiment 2, averaged over 300 (left) or 50 (right) voxels with the highest component weights. Data is from all subjects (N=8) and Texform conditions are excluded. Shares legend with B. (D) Same as C, but including Texform conditions (N=6)

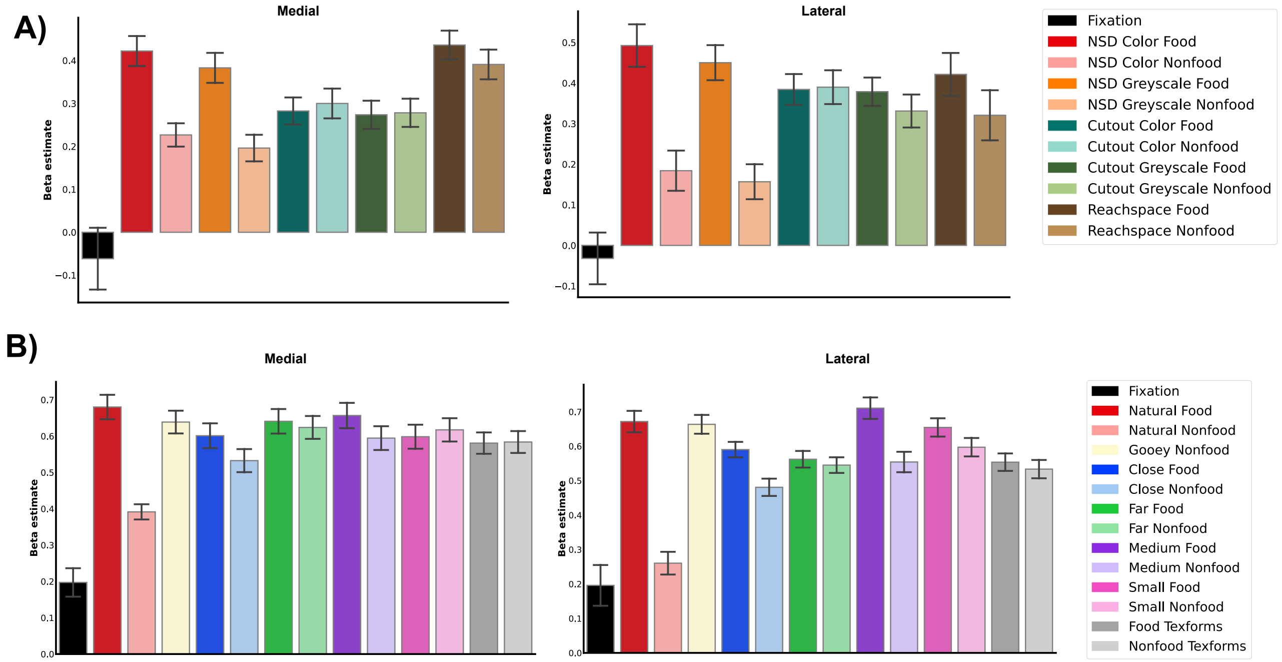


**Supplemental Fig. S5. Responses of medial and lateral bands to experiment conditions.** We used a local PCA on each individual subject and hemisphere's ventral surface to find the axis capturing medial-lateral spread and set a boundary between the two bands. We then used this partitioning on the top 300 voxels within each subject to separate medial voxels and lateral voxels. Bars show the average beta values across conditions. (A) Experiment 1 (B) Experiment 2. Plots show data from 6 subjects who saw Texform condition.

### Supplemental Section S6

#### S6A. Exploratory Analysis: Is the Food Component Response Correlated with Distance or Hand Action Affordance Depicted in NSD Images?

To explore possible effects of food salience, distance of the foreground object, distance of the background, and hand action affordance, we first collected ratings for each of these dimensions on Prolific, asking 10 participants to rate each image on the salience of that dimension on a scale from 1 to 10. For example, online participants rated “How noticeable/prominent is food in this image? (0 = no food content; 10 = extremely noticeable food content)". Intersubject reliability was significant but not high, so two of the authors also performed the same rating. Because our inter-author correlation was higher than for the participants on Prolific, we used our ratings for the main analyses, but we note that the pattern of results did not differ substantially when using the online data. These ratings were then normalized to a range between zero and 1. We then calculated the correlation between each of these factors in the stimulus set and found that all were significantly correlated with each other (Figure 4B), underlining the challenge of interpreting neural responses to natural images.

In an effort to glean what we could from the published food component responses to these images, we fit a linear model without interactions on the effect of these four factors on the magnitude of the food component response (Table S6A.i). This analysis found that each of the four factors had a significant effect, but food salience had the largest beta estimate (1.63 for food, SE = 0.10, $p < 0.001$; -0.87 for furthest distance, SE = 0.17, $p < 0.001$; -0.49 for closest distance, SE = 0.21, $p = 0.020$; 0.26 for hand actions, SE = 0.13, $p = 0.049$). The $R^2$ of this model was 0.56, meaning it explains 56% of the variance in the data. We then fit a linear model with interactions (Table S6A.ii), and again found significant effects for food salience (B = 7.03, SE = 1.22, $p < 0.001$), furthest distance (B = -2.17, SE = 0.41, $p < 0.001$), and closest distance (B = -2.58, SE = 0.56, $p < 0.001$). Hand action affordance was not significant (B = 0.24, SE = 0.37, $p = 0.526$). This analysis also found many significant multiway interactions that were hard to interpret. The $R^2$ of this model was 0.64. Finally, to account for multicollinearity, we ran a ridge regression (Table S6A.iii) and found beta values for food salience (6.43), closest distance (-2.60), furthest distance (-2.19), hand actions (0.21), along with large multiway interactions. Taken together, these exploratory analyses suggest that the food component response is strongly correlated with food salience, foreground object distance, and background distance, but not strongly affected by hand action affordance.

**Supplemental Table S6A.i. Food Salience x Distance x Hand Actions Linear Model (No Interactions) Fixed effects**

| Predictor | Estimate | SE | p-value | CI |
|---|---|---|---|---|
| Intercept | 0.24 | 0.09 | 0.006 | [0.07, 0.41] |
| Food Salience | 1.63 | 0.10 | < .001 | [1.44, 1.82] |
| Hand Actions | 0.26 | 0.13 | 0.049 | [0.00, 0.51] |
| Closest Distance | -0.49 | 0.21 | 0.020 | [-0.90, -0.08] |
| Furthest Distance | -0.87 | 0.17 | < .001 | [-1.20, -0.55] |

**Supplemental Table S6A.ii. Food Salience x Distance x Hand Actions Linear Model (With Interactions) Fixed effects**

| Predictor | Estimate | SE | p-value | CI |
|---|---|---|---|---|
| Intercept | 0.88 | 0.19 | < .001 | [0.52, 1.25] |
| Food Salience | 7.03 | 1.22 | < .001 | [4.64, 9.43] |
| Hand Actions | 0.24 | 0.37 | 0.526 | [-0.49, 0.97] |
| Closest Distance | -2.58 | 0.56 | < .001 | [-3.67, -1.48] |
| Furthest Distance | -2.17 | 0.41 | < .001 | [-2.97, -1.36] |
| Food Salience x Hand Actions | -6.48 | 1.49 | < .001 | [-9.40, -3.55] |
| Food Salience x Closest Distance | -27.99 | 6.12 | < .001 | [-39.99, -15.98] |
| Hand Actions x Closest Distance | -2.05 | 2.00 | 0.304 | [-5.97, 1.86] |
| Food x Furthest Distance | -24.67 | 5.14 | < .001 | [-34.75, -14.59] |
| Hand Actions x Furthest Distance | 0.28 | 1.45 | 0.847 | [-2.57, 3.13] |
| Closest Distance x Furthest Distance | 4.45 | 1.02 | < .001 | [2.45, 6.45] |
| Food x Hand x Closest | 37.38 | 8.22 | < .001 | [21.25, 53.51] |
| Food x Hand x Furthest | 26.55 | 6.80 | < .001 | [13.22, 39.89] |
| Food x Closest x Furthest | 88.74 | 16.18 | < .001 | [56.98, 120.50] |
| Hand x Closest x Furthest | -0.96 | 5.82 | 0.869 | [-12.38, 10.46] |
| Food x Hand x Closest x Furthest | -105.22 | 22.99 | < .001 | [-150.35, -60.08] |

**Supplemental Table S6A.iii. Food Salience x Distance x Hand Actions Ridge Regression (With Interactions)**

| Predictor | Beta |
|---|---|
| Food Salience | 6.43 |
| Hand Actions | 0.21 |
| Closest Distance | -2.60 |
| Furthest Distance | -2.19 |
| Food Salience x Hand Actions | -5.75 |
| Food Salience x Closest Distance | -26.12 |
| Food Salience x Furthest Distance | -22.13 |
| Hand Actions x Closest Distance | -1.94 |
| Hand Actions x Furthest Distance | 0.37 |
| Closest Distance x Furthest Distance | 4.51 |

| Predictor | Beta |
|---|---|
| Food x Hand x Closest | 35.34 |
| Food x Hand x Furthest | 23.45 |
| Food x Closest x Furthest | 81.57 |
| Hand x Closest x Furthest | -1.32 |
| Food x Hand x Close x Far | -97.44 |

**S6B: Food CNN Models**

Due to the confounding of multiple image properties in NSD images (see Figure 4A), we sought to use new, controlled stimuli to specifically test certain hypotheses. We used a CNN model of the food component reported in Khosla et al (2022), the predictions of which had a 0.83 correlation with the actual component response on held-out NSD data. Given the high predictivity of this CNN model of the food component (r = 0.83 with held-out NSD data, r = 0.63 with Experiment 1, r = 0.57 with Experiment 2), we decided to pilot novel stimuli on the model to predict the component response to these stimuli. Findings are shown below.

**(i) Different sized scenes:** In the NSD shared 1000 images, we see some effect of distance (Figure 4). We therefore decided to run exploratory tests on whether the food component model response depends on the size and viewing distance of a scene. We used the Scene Categories by Size dataset (Park et al 2015), which contains six categories of scenes ranging in sizes with D1 being the closest (closets, bathrooms) and D6 being the furthest (airports, stadiums). The predictions of the CNN-based food component model to all categories D1-D6 were strikingly low, near the model's predictions to NSD Non-food. Although the predicted response decreases with distance, the effect of distance seems to plateau as the scene size increases. We fit a regression model to only the D1-D6 conditions of distance as a predictor of predicted response, and found that a quadratic model provided a significantly better fit than a linear model ($\Delta R^2 = 0.19$, $F(1, 285) = 86.42$, $p < 0.001$), explaining 39% of the variance in the data ($R^2 = 0.39$) We compared the responses of D1 and D5, which respectively had the highest and lowest predicted component response, to NSD non-food, and found that the model predicted a significantly lower response to NSD non-food than either of these categories (NSD Non-food – D1: estimate = -5.35e-05, 95% CI = [-6.43E-05, -4.27E-05], p < 0.001; NSD Non-food – D5 estimate = -2.55E-05, 95% CI = [-3.64E-05, -1.46E-05], p < 0.001). Altogether these results indicate that the CNN-based food component model predicts a significant, but non-linear, effect of viewing distance on component response. Additionally, there is a large difference in predicted response to NSD food and the closest distance category D1, indicating that either the absence of food information, or the presence of scene information, even at the viewing distance of a small closet or shower, may drive a large portion of the component's response.

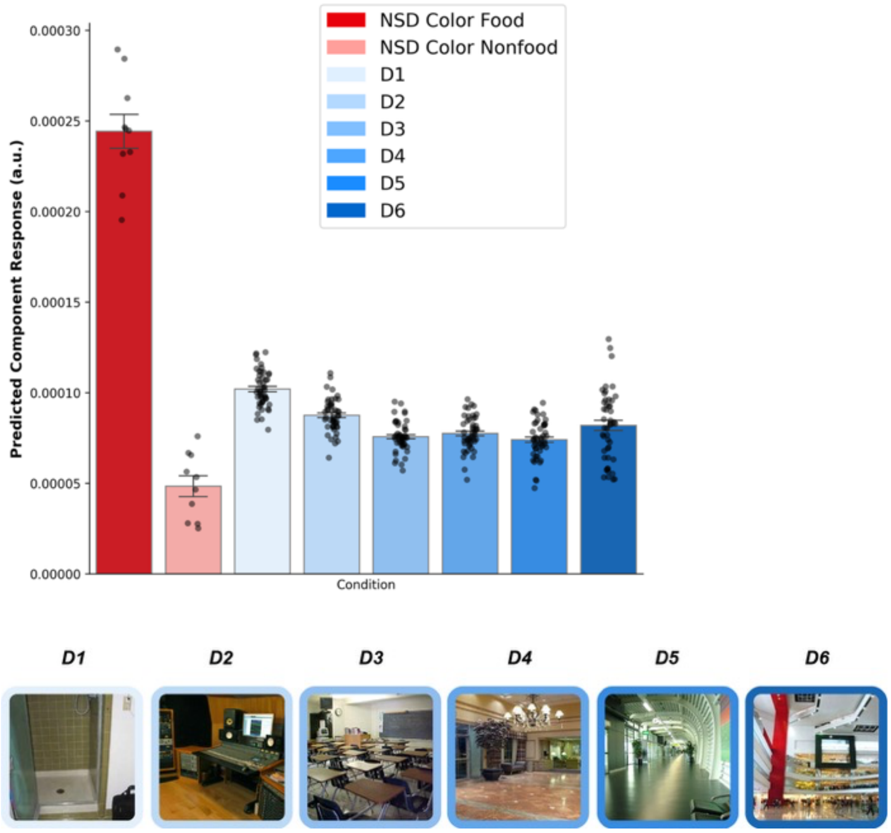


**(ii) Mid-distances with and without food:** This image set features images of indoor mid-distances (about 10-20 ft), with and without a pizza in the middle. The pizza is placed in a noticeable position in the center of the frame, about 10-15 feet away from the camera. The responses to the mid-distance images both with and without food were strikingly low, suggesting that the presence of scene information, or the increased distance of food from the camera, drives a large portion of the component response. Additionally, the model did not predict a significant difference in response between the mid distance images with and without food (estimate = 4.30E-07, 95% CI [-1.88E-05, 1.96E-05], p = 0.964). This motivated the idea that the component may not be selective for food when the food is too far away, which we further tested in Experiment 2.

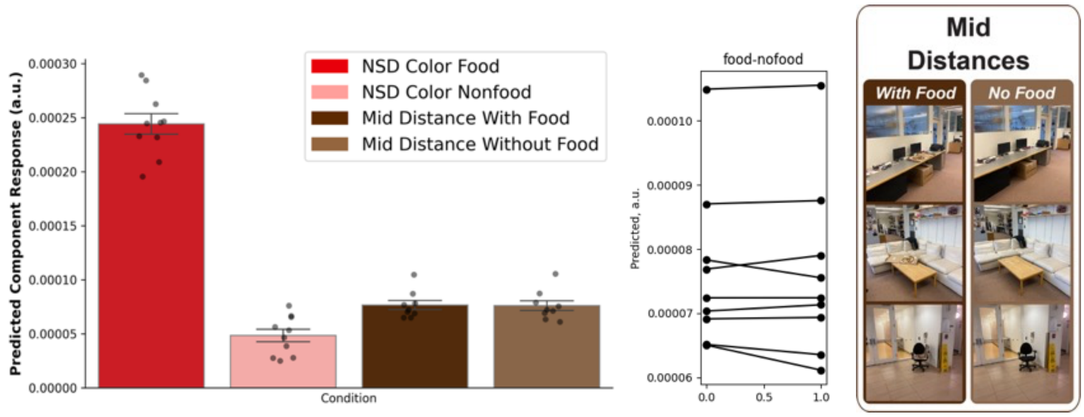

**(iii) RW size & animacy:** We next ran exploratory tests of real-world size because a) differential responses to small and large objects (and animate/inanimate objects) have been reported before in the ventral pathway (Konkle & Caramazza, 2013) and b) in NSD images, food tends to be of a mid-small size, like a plate, while non-food images with lowest component response depicted larger objects like cars and horses. In NSD data, real-world size is confounded with other features such as distance and scene information, and thus we tested new stimuli on the CNN-based food component model to isolate impacts of RW size and animacy. For this analysis we used the RW size x animacy dataset (Konkle & Caramazza, 2013). We found that the CNN food model predicts a higher mean response to small over big (estimate = 3.25E-05, 95% CI [2.79E-05, 3.71E-05], $p < 0.001$), a lower mean response to animate than inanimate (estimate = -1.33E-05, 95% CI [-1.79E-05, -8.68E-06], $p < 0.001$), but no significant interaction (estimate = -7.82E-06, 95% CI [-1.71E-05, 1.42E-06], $p = 0.097$). Based on these exploratory findings, we decided  to test RW size in Experiment 2.

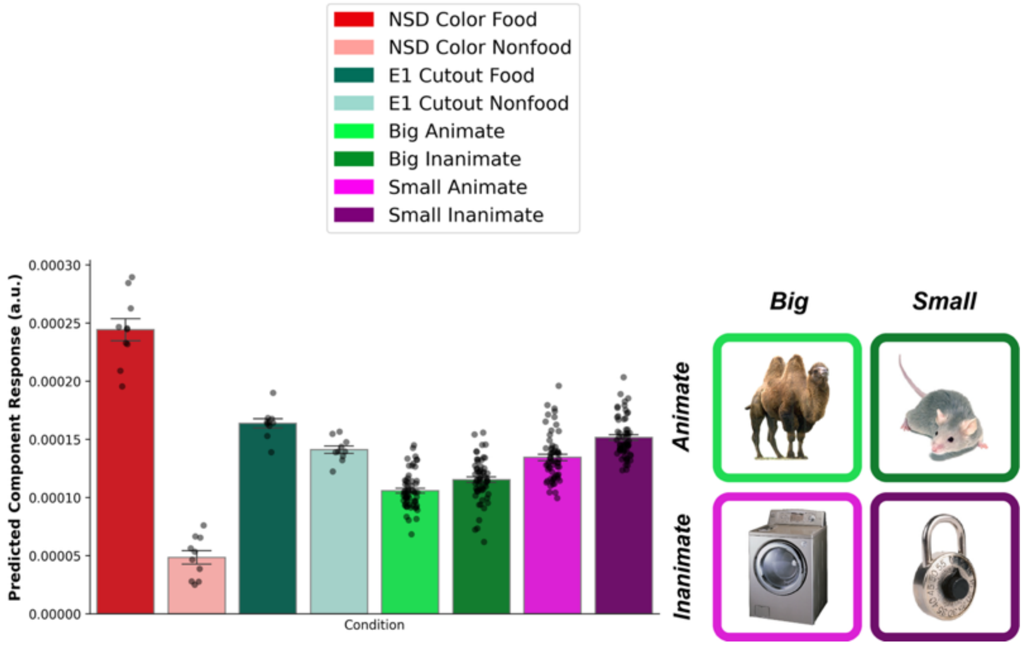


**(iv) Hard food vs. gooey non-food:** Food tends to have a gooey/soft material property, which can be seen in the NSD food we included in our Experiment 1. Might the food component response be selective for this material property, rather than food per se? To test this idea on our CNN-based food model, we measured responses to hard food (hard candies, pretzels, etc.) with gooey non-foods (slime, Tide pods). If the component is more selective for the material property of softness/gooeyness than foodness, we would expect a higher response to gooey non-food than to hard food. Although the predicted response to gooey non-food was smaller than that of hard food (estimate = 3.06E-05, 95% CI [7.38E-06, 5.39E-05], $p = 0.011$), we did find a significant interaction such that the difference between hard food and gooey non-food was smaller than the difference between NSD Food and NSD Non-food (estimate = 1.65E-04, 95% CI [1.31E-04, 1.99E-04], $p < 0.001$), and the predicted response to gooey non-food was significantly higher than NSD non-food (estimate = -1.47E-04, 95% CI [-1.72E-04, -1.23E-04], $p < 0.001$). Due to the surprisingly high predicted response to gooey non-food, we decided to test the material property hypothesis in new fMRI subjects in Experiment 2.

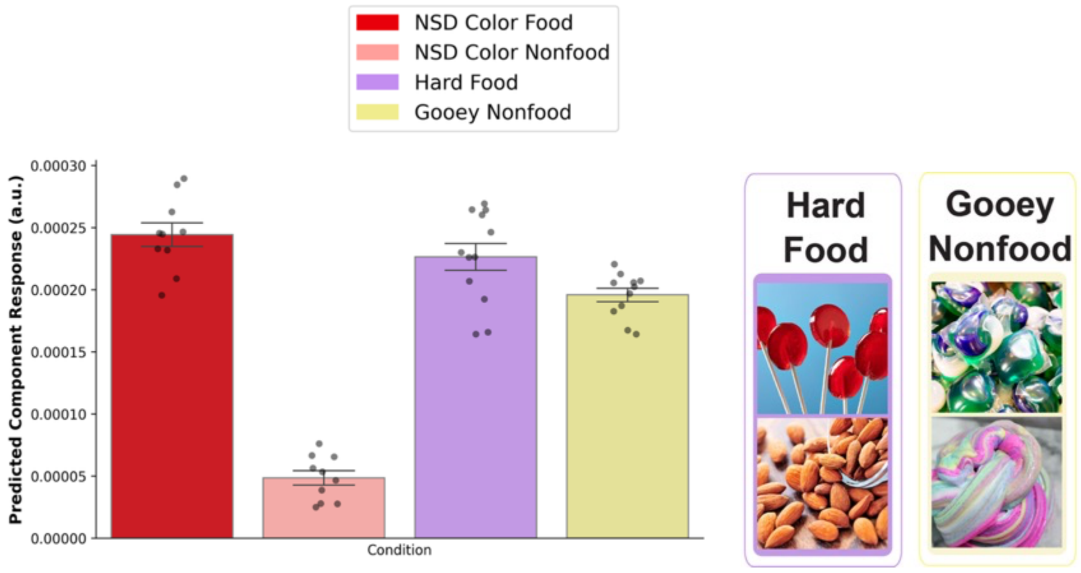


**(v) Food Context with and without food:**
To test the role of food contexts, we created stimuli featuring a 2x2 design of food and non-food contexts with and without food. Photographs were taken in-house. Food contexts included bowls, tables, plates, and to-go containers. Non-food contexts included sidewalks, stairs, and floors. Food context had a significantly higher predicted response than non-food context within both food images (estimate = 1.95E-05, 95% CI [2.13E-06, 3.68E-05], p = 0.029) and non-food images (estimate = 1.87E-05, 95% CI = [1.38E-06, 3.60E-05], p = 0.035), and there was a significant effect of food within both food contexts (estimate = 2.41E-05, 95% CI = [6.76E-06, 4.14E-05], p = 0.007) and non-food contexts (estimate = 2.33E-05, 95% CI = [6.01E-06, 4.07E-05], p = 0.009). There was no significant difference between the effect of food within food context or non-food context (estimate = 7.45E-07, 95% CI = [-2.87E-05, 3.02E-05], p = 0.998). The effect of food was significantly greater between NSD food and NSD non-food than within both food context (interaction, estimate = -1.72E-04, 95% CI = [-1.96E-04, -1.47E-04], p < 0.001), and non-food context (interaction, estimate = -1.72E-04, 95% CI = [-1.97E-04, -1.47E-04], p < 0.001). These results show that while the food and non-food contexts tested here have some effect on the predicted response, the effect size was small and could not account for the high response to NSD food or the low response to NSD non-food.

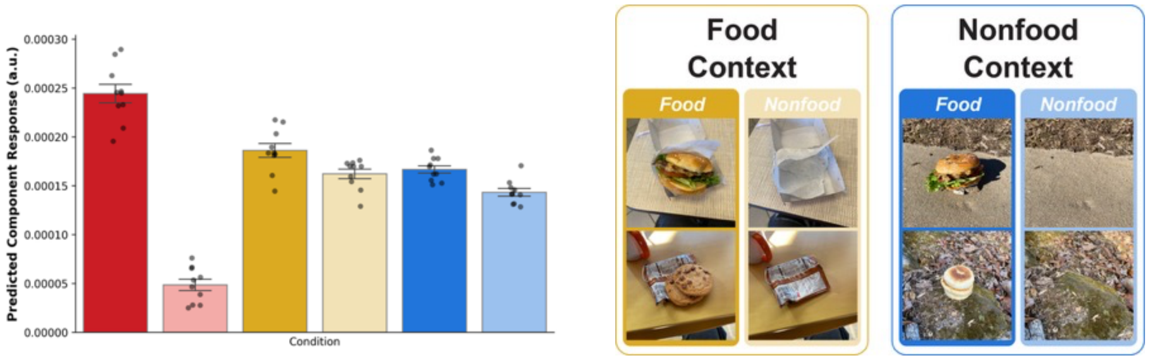


**(vi) Color Change:** Does the food component 'dislike' colors that are not common for food? Behaviorally, we may be hesitant to eat something like spaghetti with blue sauce, because it is unclear if it is edible or not. Although Experiment 1 showed a significant effect of color over greyscale images, it was unknown whether the component response is impacted by the hue, rather than just the presence of color. We altered NSD food images, changing the colors to hues uncommon in food, such as blue and purple. The color-changed food images had a significantly lower response than NSD food images (estimate = 3.52E-05, 95% CI = [1.23E-05, 5.81E-05], p = 0.004), but were not significantly different from greyscale food (estimate = 5.77E-06, 95% CI = [-1.71E-05, 2.87E-05], p = 0.610).

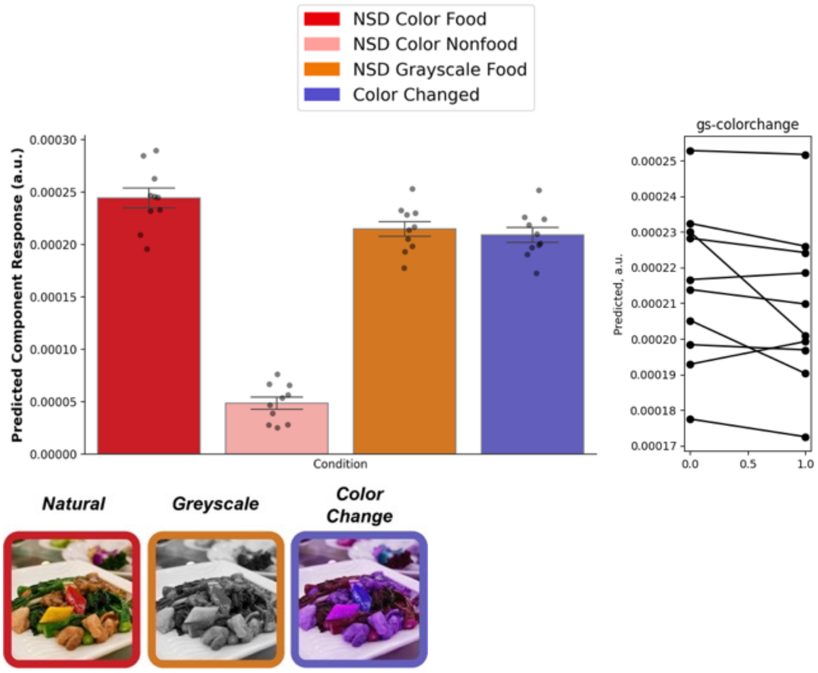

**Supplemental Section S7. PCA Analysis of NSD stimuli**

To gain some insight into the low-level features or image statistics that differentiate food from other categories, we passed the 1000 shared NSD images through AlexNet, and then performed PCA on responses of units in layers conv1, conv2, conv3, and fc6. We also performed PCA directly on the image pixels. While food images did not cluster in the PC space of the image pixels, they did cluster in AlexNet fc6, with emergence of these groups in the conv2 layer. The principal components were not fully interpretable, but in the conv3 layer, PC2 appears to correspond to food, along with PC3 in the fc6 layer. PC plots, as well as the top and bottom image of each of the corresponding PCs, are plotted below. These analyses were exploratory and not preregistered. Photographs of people have been redacted.

**Pixel space**

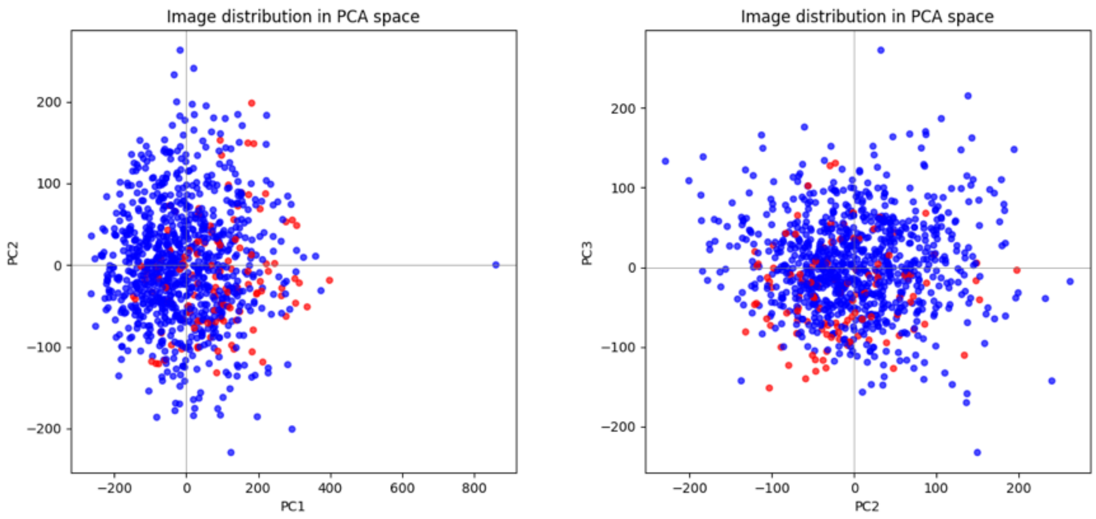


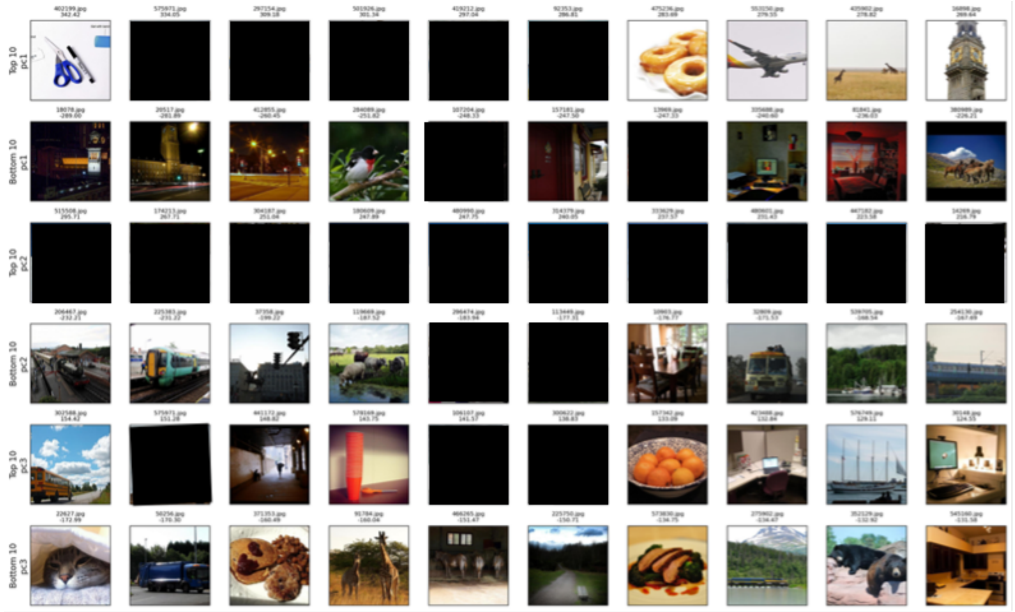

### Conv1

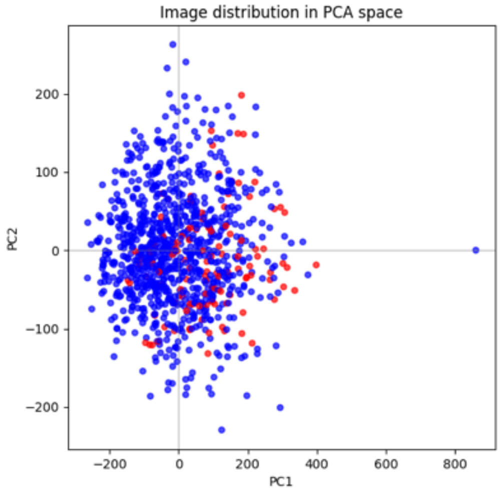
Image distribution in PCA space
PC1
PC2

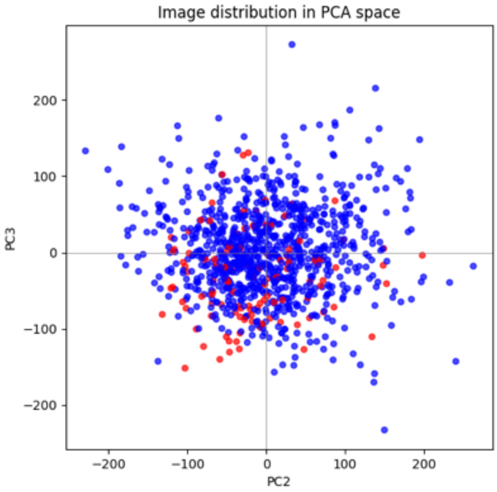
Image distribution in PCA space
PC2
PC3

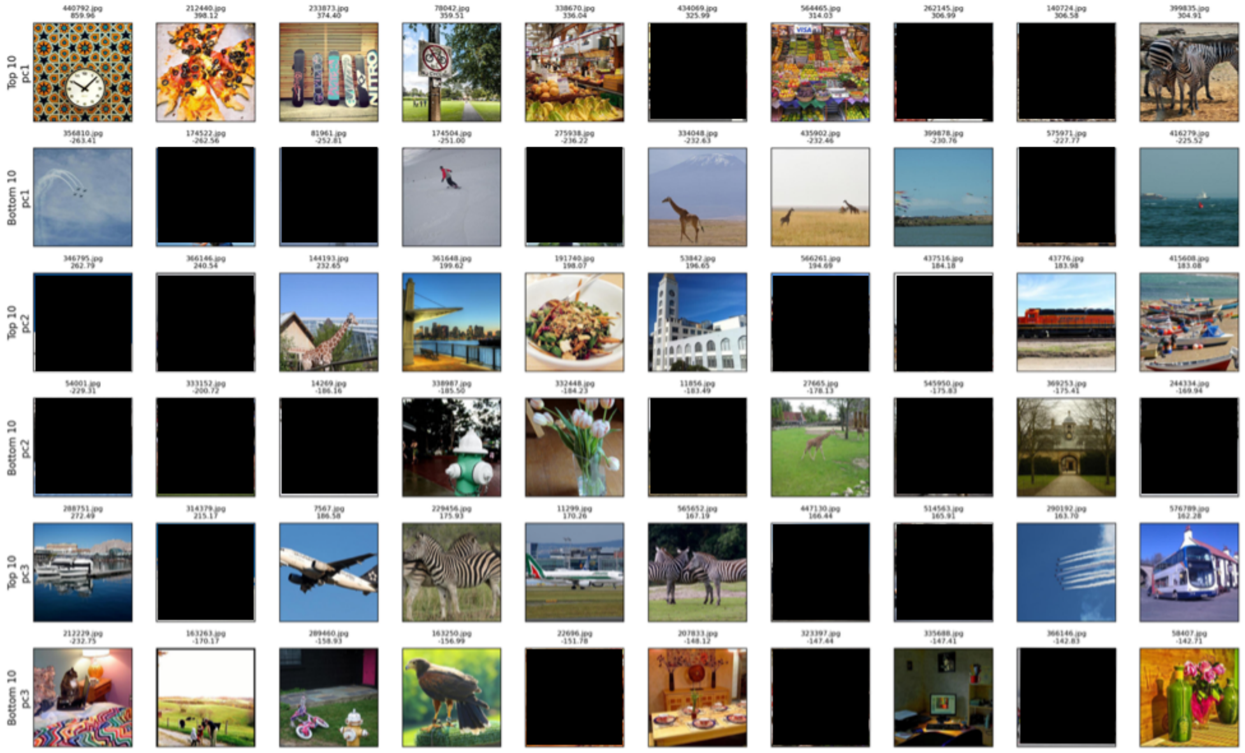
Top 10 pc1
Bottom 10 pc1
Top 10 pc2
Bottom 10 pc2
Top 10 pc3
Bottom 10 pc3

### Conv2

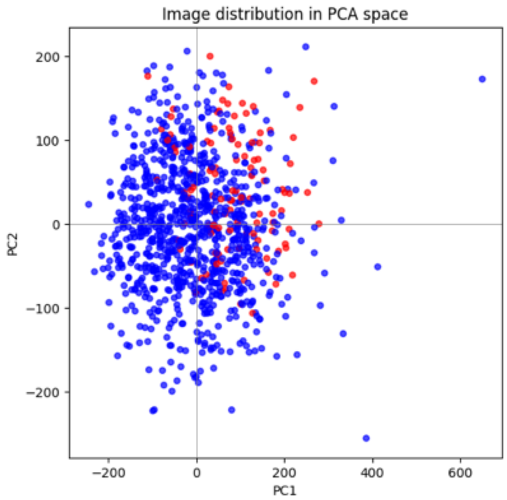
Image distribution in PCA space
PC1
PC2

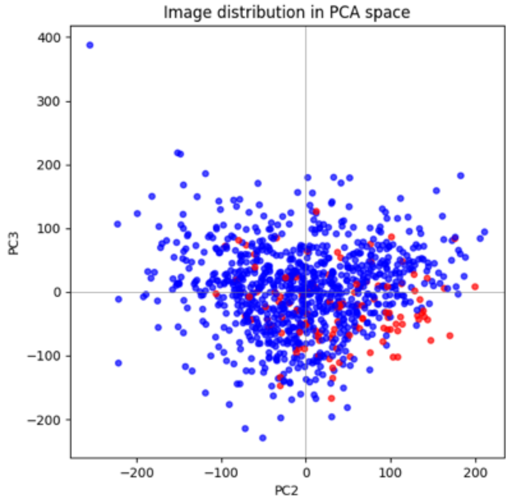
Image distribution in PCA space
PC2
PC3

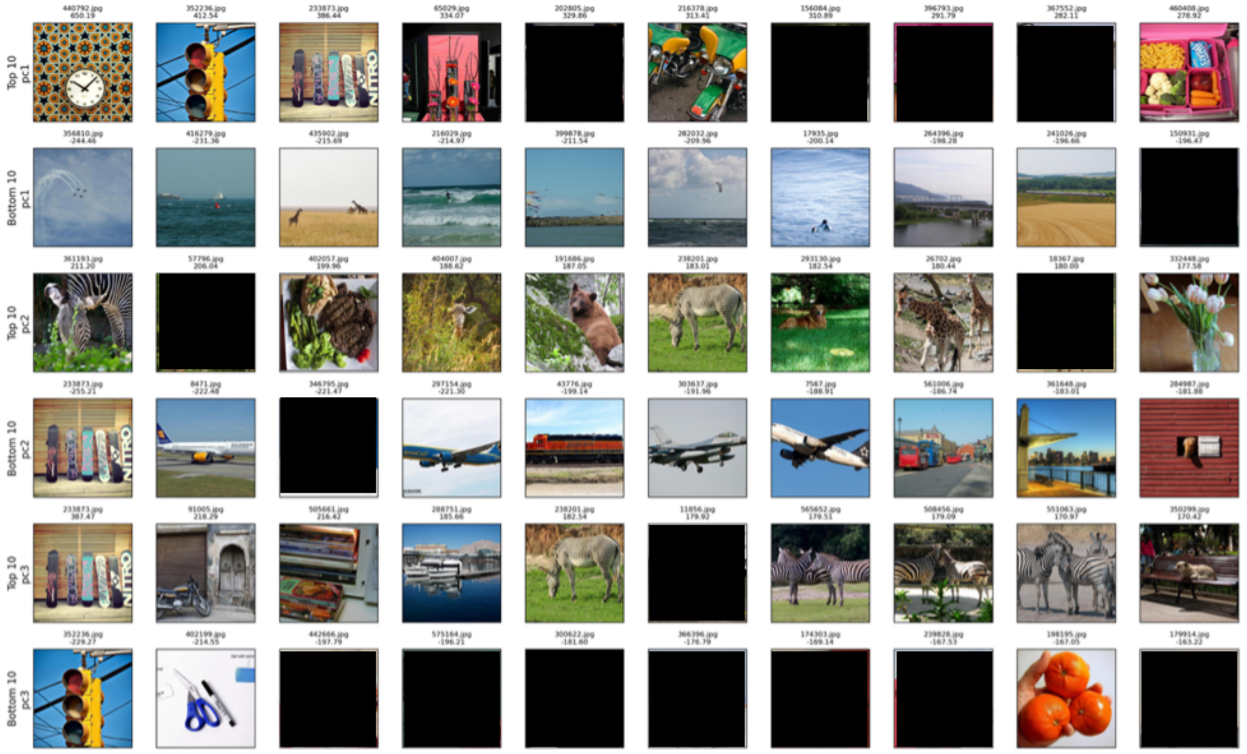
Top 10 pc1
Bottom 10 pc1
Top 10 pc2
Bottom 10 pc2
Top 10 pc3
Bottom 10 pc3

### Conv3

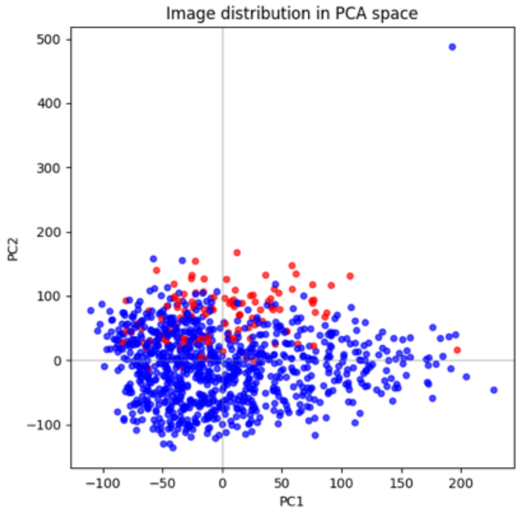
Image distribution in PCA space
PC1
PC2
−100
−50
0
50
100
150
200
300
400
500

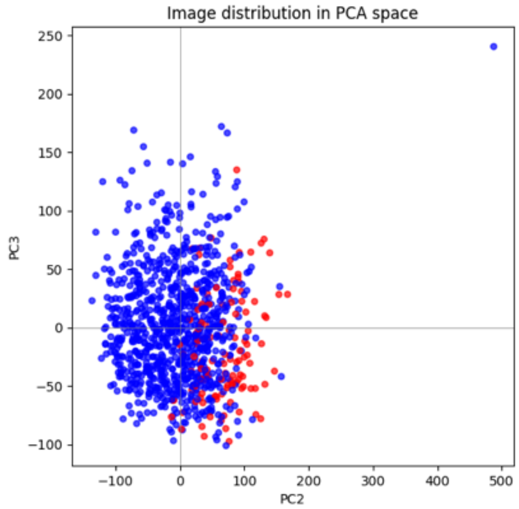
Image distribution in PCA space
PC2
PC3
−100
−50
0
50
100
150
200
250
300
400
500

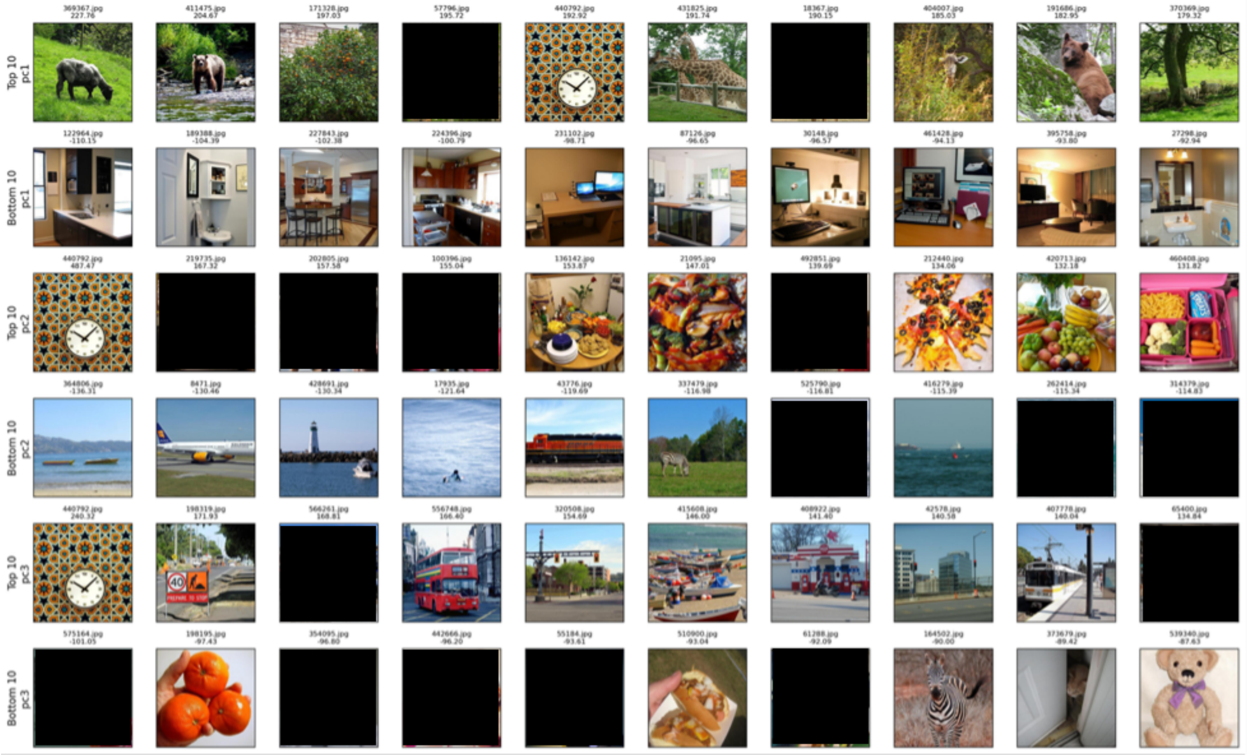
Top 10 pc1
Bottom 10 pc1
Top 10 pc2
Bottom 10 pc2
Top 10 pc3
Bottom 10 pc3

## Fc6

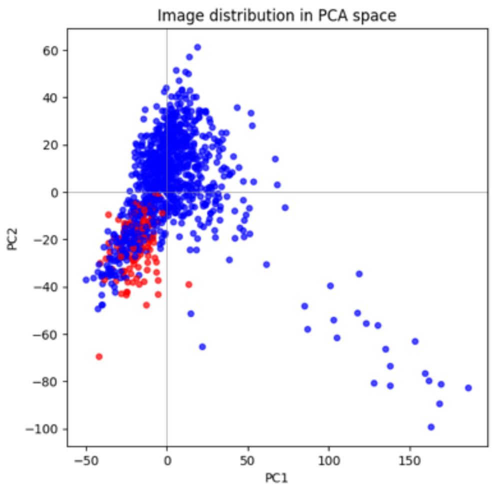

Image distribution in PCA space
PC1
PC2


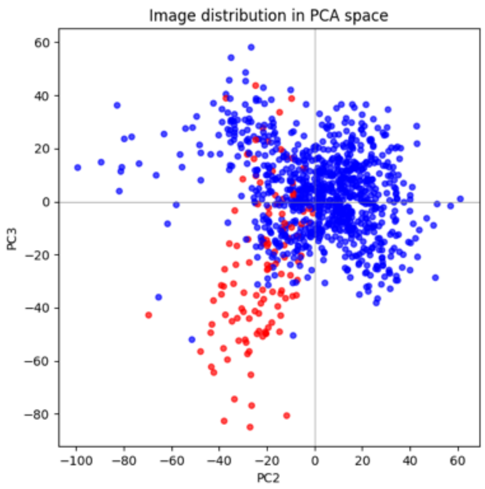

Image distribution in PCA space
PC2
PC3


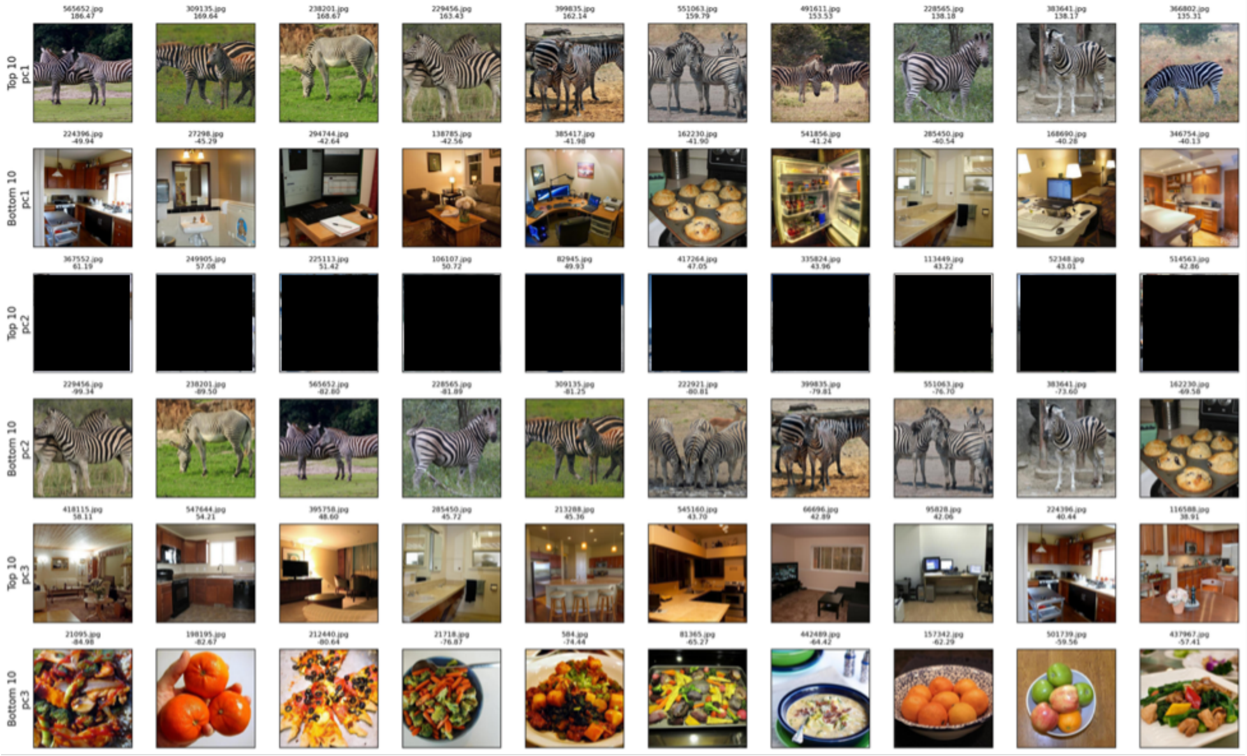

Top 10 pc1
Bottom 10 pc1
Top 10 pc2
Bottom 10 pc2
Top 10 pc3
Bottom 10 pc3

**Supplemental Section S8. RT Experiment**

We hypothesize that if there are visual statistics that are common in food, and that the "food component" is responsive to these visual statistics, it might function as a first-level filter to detect whether the visual input is likely to be food, before sending the information to a later processing stage that might more definitively detect or analyze food. If this were the case, then we would expect it to take longer to recognize non-food as non-food if it has a high food component response, and conversely it would take longer to recognize food as food if it has a low food component response. We tested this using an online behavioral experiment in which we recruited participants on Prolific (N=6). Participants were shown each of the 1000 shared NSD images, for 300ms each, and they were asked to make a judgment of whether it was food or non-food as fast as possible. We found trends as hypothesized such that there was a negative correlation between RT and component response in food images ($r = -0.48$, $p < 0.001$) and a positive correlation between RT and component response in non-food images ($r = 0.38$, $p < 0.001$). We also collected RTs for our images from Experiments 1 and 2, but did not see clear trends of correlation between RT and component response. (See Fig. S8) Together, these results suggest that the food component response captures visual features that facilitate food categorization in naturalistic images, but that this relationship between RT and component response does not generalize to out-of-distribution images in which many of these visual features are reduced or removed.

*NSD Shared 100 Images*

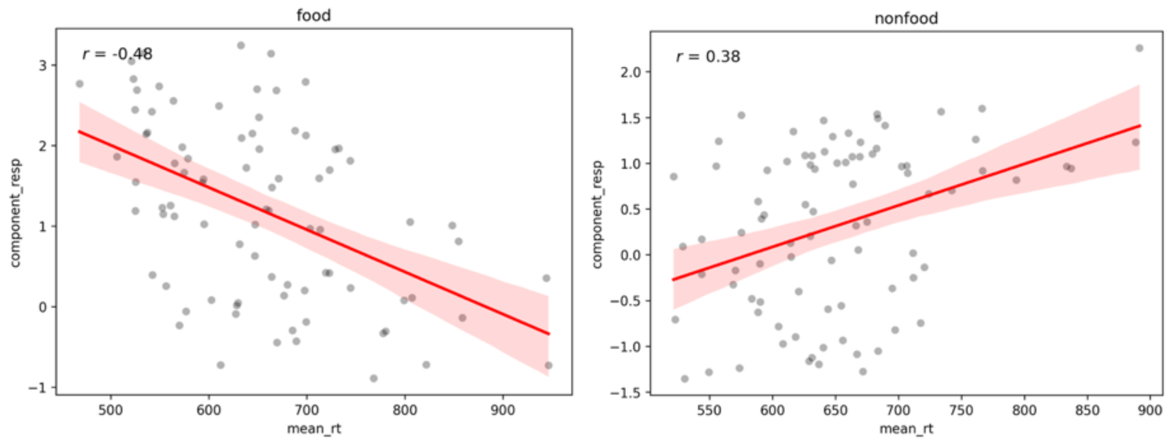


*Experiment 1 Stimuli*

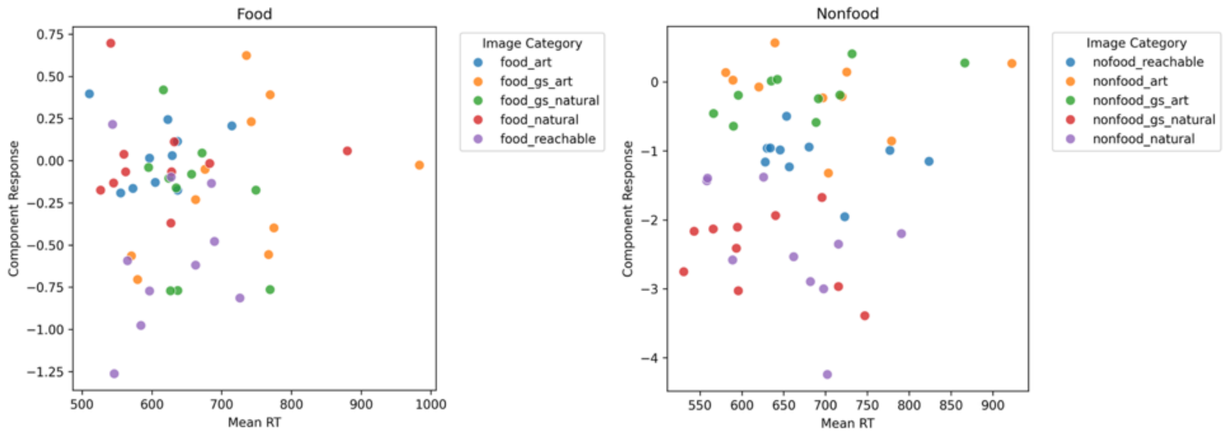


*Experiment 2 Stimuli*

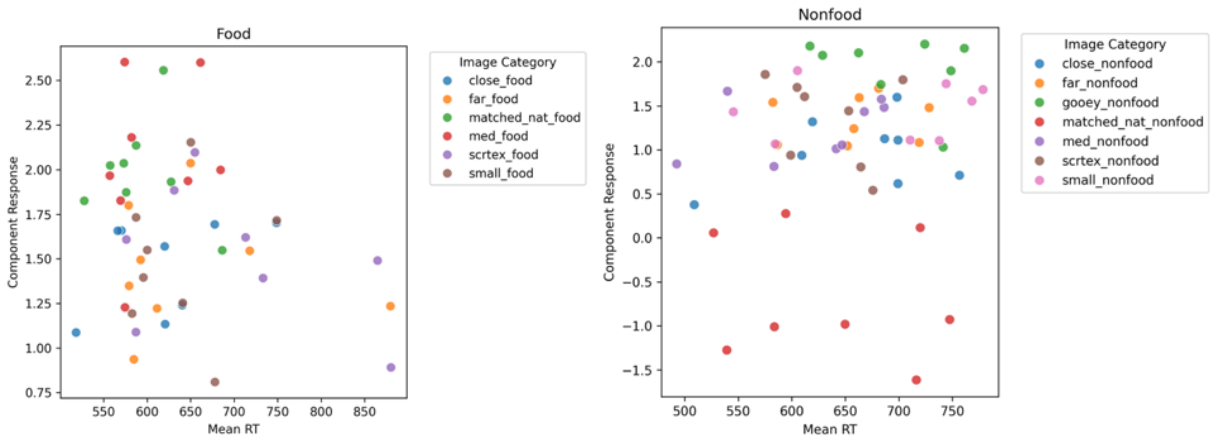


**Supplemental Fig. S8.** The first row shows the relationship between RT for determining whether the image is food or non-food and component response in NSD subjects to 1000 shared NSD images, for food (left) and non-food (right) separately. The second row shows the same for images from Experiment 1, color coded by category. The third row shows the same for images from Experiment 2, color coded by category.

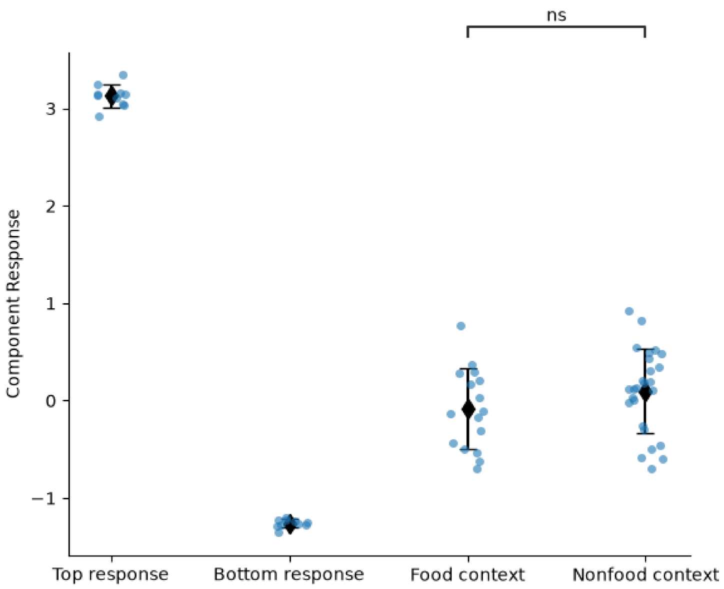


**Supplemental Fig. S9.** Average NSD component response to images with food contexts and images with non-food contexts when there is no food in the image. The food contexts are images of tabletops and kitchens without food. Non-food contexts consist largely of bathrooms, in order to approximately match semantic and low-level visual features of the food context images. In order to contextualize the response magnitude within the range of all images, “Top response” shows the images with the top NSD component response and “Bottom response” shows images with the bottom NSD component response. Each point shows one image and its component response averaged across NSD subjects.
